# The genome of the jellyfish *Clytia hemisphaerica* and the evolution of the cnidarian life-cycle

**DOI:** 10.1101/369959

**Authors:** Lucas Leclère, Coralie Horin, Sandra Chevalier, Pascal Lapébie, Philippe Dru, Sophie Peron, Muriel Jager, Thomas Condamine, Karen Pottin, Séverine Romano, Julia Steger, Chiara Sinigaglia, Carine Barreau, Gonzalo Quiroga Artigas, Antonella Ruggiero, Cécile Fourrage, Johanna E. M. Kraus, Julie Poulain, Jean-Marc Aury, Patrick Wincker, Eric Quéinnec, Ulrich Technau, Michaël Manuel, Tsuyoshi Momose, Evelyn Houliston, Richard R. Copley

**Author notes:** Current addresses: P.L. Architecture et Fonction des Macromolécules Biologiques, Aix-Marseille Université, Marseille, France; J.S.: Department for Molecular Evolution and Development, Centre of Organismal Systems Biology, University of Vienna, Vienna A-1090, Austria; C.S. Institut de Génomique Fonctionnelle de Lyon (IGFL), École Normale Supérieure de Lyon, CNRS UMR 5242 - INRA USC 1370, 69364 Lyon cedex 07, France; G.Q.A.:The Whitney Laboratory for Marine Bioscience, University of Florida, St. Augustine, FL, 32080, USA; A.R. Centre de Recherche de Biologie cellulaire de Montpellier (CRBM), CNRS UMR 5237, Université de Montpellier, 1919 Route de Mende, 34293 Montpellier Cedex 5, France; C.F. Service de Génétique UMR 781, Hôpital Necker - APHP, Paris, France; J.E.M.K. Sars International Centre for Marine Molecular Biology, University of Bergen, Thormøhlensgate 55, 5006 Bergen, Norway.

## Abstract

Jellyfish (medusae) are a distinctive life-cycle stage of medusozoan cnidarians. They are major marine predators, with integrated neurosensory, muscular and organ systems. The genetic foundations of this complex form are largely unknown. We report the draft genome of the hydrozoan jellyfish *Clytia hemisphaerica* and use multiple transcriptomes to determine gene use across life-cycle stages. Medusa, planula larva and polyp are each characterised by distinct transcriptome signatures reflecting abrupt life cycle transitions, and all deploy a mixture of phylogenetically old and new genes. Medusa specific transcription factors, including many with bilaterian orthologs, associate with diverse neurosensory structures. Compared to *Clytia*, the polyp-only hydrozoan *Hydra* has lost many of the medusa-expressed transcription factors, despite similar overall rates of gene content and sequence evolution. Absence of expression and gene loss among *Clytia* orthologs of genes patterning the anthozoan aboral pole, secondary axis and endomesoderm support simplification of planulae and polyps in Hydrozoa, including loss of bilateral symmetry. Consequently, although the polyp and planula are generally considered the ancestral cnidarian forms, in *Clytia* the medusa maximally deploys ancestral cnidarian–bilaterian transcription factor gene complexity.

---

In most cnidarians a ciliated, worm-like planula larva settles to produce a polyp. In Anthozoa (corals and anemones), the polyp is the sexually reproductive form, but in the Medusozoa branch of Cnidaria, polyps generally produce sexually reproductive jellyfish by a process of strobilation or budding. Jellyfish (medusae) are gelatinous, pelagic, radially symmetric forms found only in the medusozoans. They show complex physiology and behaviour based on neural integration of well-defined reproductive organs, digestive systems, locomotory striated muscles and sensory structures, even camera eyes in some species. Medusae in many species show some degree of nervous system condensation, notably the nerve rings running around the bell margin [1]. Historical arguments that the medusa is the ancestral cnidarian form are no longer widely supported, as recent molecular phylogenies suggest Anthozoa and Medusozoa are sister groups, favouring a benthic, polyp-like adult ancestor on parsimony grounds [2,3] (supplementary note 1). Candidate gene expression studies have shown parallels between medusa and polyp development [4], and transcriptome comparisons between species with and without medusae have extended candidate gene lists [5], but in general the genetic foundations of complex medusa evolution within the cnidarian lineage are not well understood.

There are four classes of Medusozoa: the Cubozoa (box jellyfish), Scyphozoa (so-called ‘true’ jellyfish), Staurozoa (stalked ‘jellyfish’) and Hydrozoa [2,6]. Life cycles in different medusozoan lineages have undergone frequent modifications, including loss of polyp, planula and medusa stages. *Hydra*, the classical model of animal regeneration, is a hydrozoan characterized by the loss of the planula and medusa stages from the life-cycle. Compared to anthozoan genomes [7–9], the *Hydra* genome is highly diverged and dynamic; it may therefore be untypical of the medusozoa and even the Hydrozoa [10]. Here we describe the genome of *Clytia hemisphaerica*, a hydrozoan with a typical medusozoan life-cycle, including planula, polyp and medusa stages (Fig. 1). *Clytia* is easy to maintain and manipulate, and amenable to gene function analysis including Cas9 mediated mutation, allowing mechanistic insight into cellular and developmental processes [6,11,12]. We analyse transcriptomes from all life cycle forms, illuminating the evolution of the planula, polyp and medusa, and demonstrate how the gene complement of the cnidarian–bilaterian ancestor provided the foundation of anatomical complexity in the medusa.

**Figure 1.**
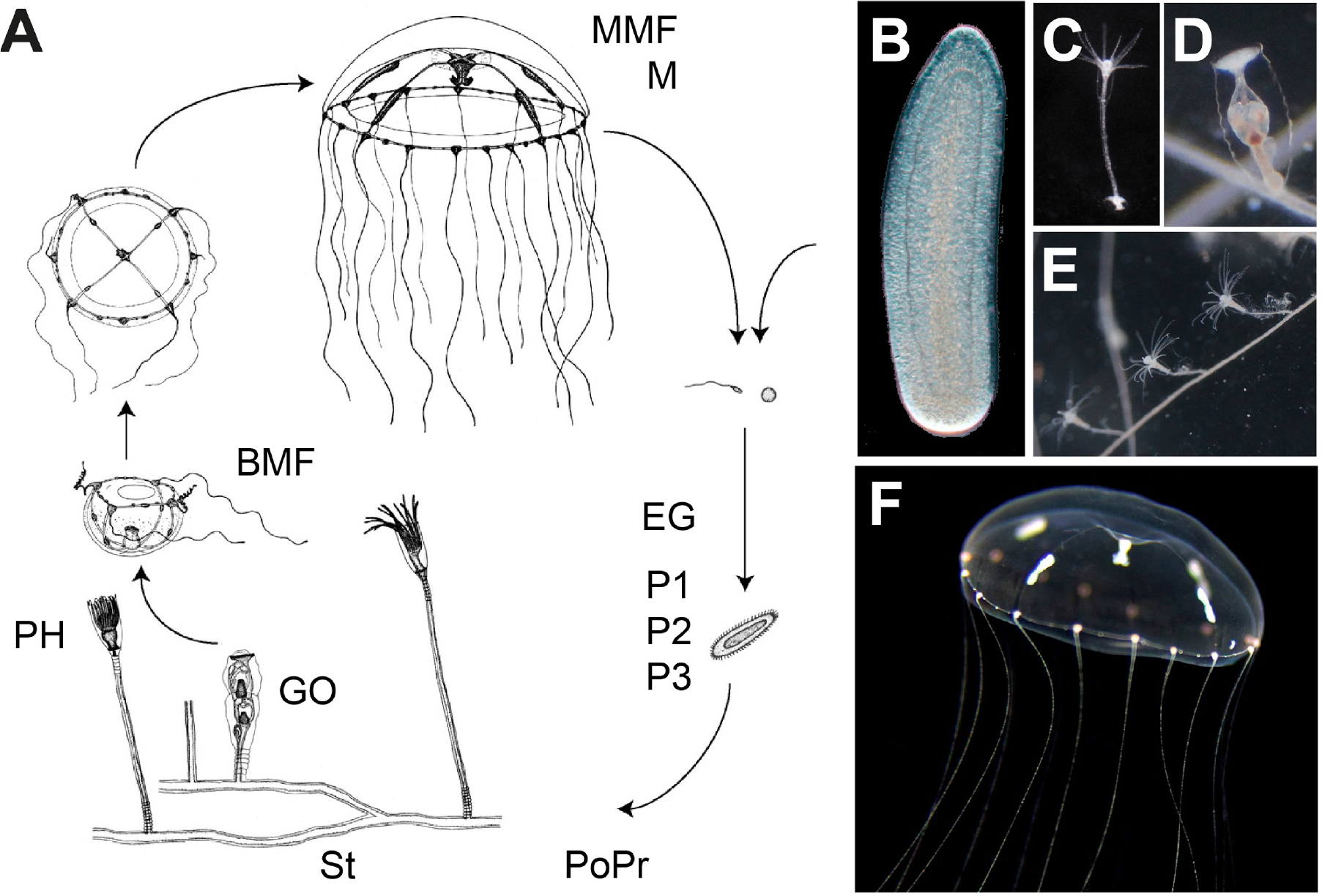
The *Clytia* life cycle (A). The planula larva (B) develops from a fertilized egg, and metamorphoses into a primary polyp (C). The polyp then extends asexually forming a colony composed of (D) feeding polyps (gastrozooids) attached through a common stolon, and gonozooids (E) that release swimming medusae (F). Abbreviations in (A) correspond to the mRNA libraries in tab. S3: EG: early gastrula; P1/P2/P3: Planula 24h-post-fertilization/48hpf/72hpf; PoPr: Primary polyp; St: Stolon: GO: Gonozooid: PH: Gastrozooid/Polyp head; BMF: Baby medusae 1d old: MMF: Mature female medusa; M: Mature Male Medusa. Drawings in (A) from [69] (F) adapted from [11].

## Results

### Characteristics of the *Clytia* genome

We sequenced the *Clytia hemisphaerica* genome using a whole genome shotgun approach (see methods, tab. S1, fig. S1), giving an assembly with overall length of 445Mb. Staining of DNA in prophase oocytes shows the genome is packaged into 15 chromosome pairs (fig. S2). We predicted transcript sequences using expressed sequence reads from a comprehensive set of stages and tissues as well as deep sequencing of mixed-stage libraries (tab. 1) giving 26,727 genes and 69,083 transcripts. BUSCO analysis of the presence of ‘Universal Single Copy Orthologs’ indicates a coverage of 86%, comparable or better than many other recently sequenced non-bilaterian animals (tab. S1). The GC content is 35%, higher than Hydra (29%, [10]) but lower than the anthozoan *Nematostella* (39%, [7]), with a repeat content of ~41%. Reads mapped to the genome suggested a polymorphism frequency of ~0.9%, likely an underestimate of heterozygosity in wild populations, as genomic DNA and mRNA for transcriptomes was derived from self-crossed laboratory-reared *Clytia* Z strains - see methods. The complete mitochondrial genome showed the same gene order as the Hydroidolina ancestor [13] (fig. S1).

### Patterns of gene gain and loss

We identified groups of orthologs for a selection of animals with completely sequenced genomes, and unicellular eukaryotic outgroups (see methods). A binary matrix of orthologous group presence or absence was then used to infer a maximum-likelihood phylogeny that recapitulated the widely accepted major groupings of animals (fig. 2); Cnidarians were the sister group of the Bilateria, and within the Cnidaria, we recovered the expected monophyletic relationships: corals, anemones, anthozoans and hydrozoans. The hydrozoan branch lengths were the longest within the cnidarians, implying elevated rates of gene gain and loss in their lineage, although branches leading to several other species were noticeably longer, including the ecdysozoan models *Caenorhabditis* and *Drosophila*, the ascidian *Ciona*, as well as the ctenophore *Mnemiopsis*. *Clytia* and *Hydra* branch lengths were similar, suggesting that genome evolution has proceeded at comparable rates in these two hydrozoan lineages. This gene content-based phylogeny positioned sponges (represented by *Amphimedon*), not ctenophores (represented by *Mnemiopsis*), as the sister group of all other animals [14–16], although confirmation with additional species is required.

**Figure 2.**
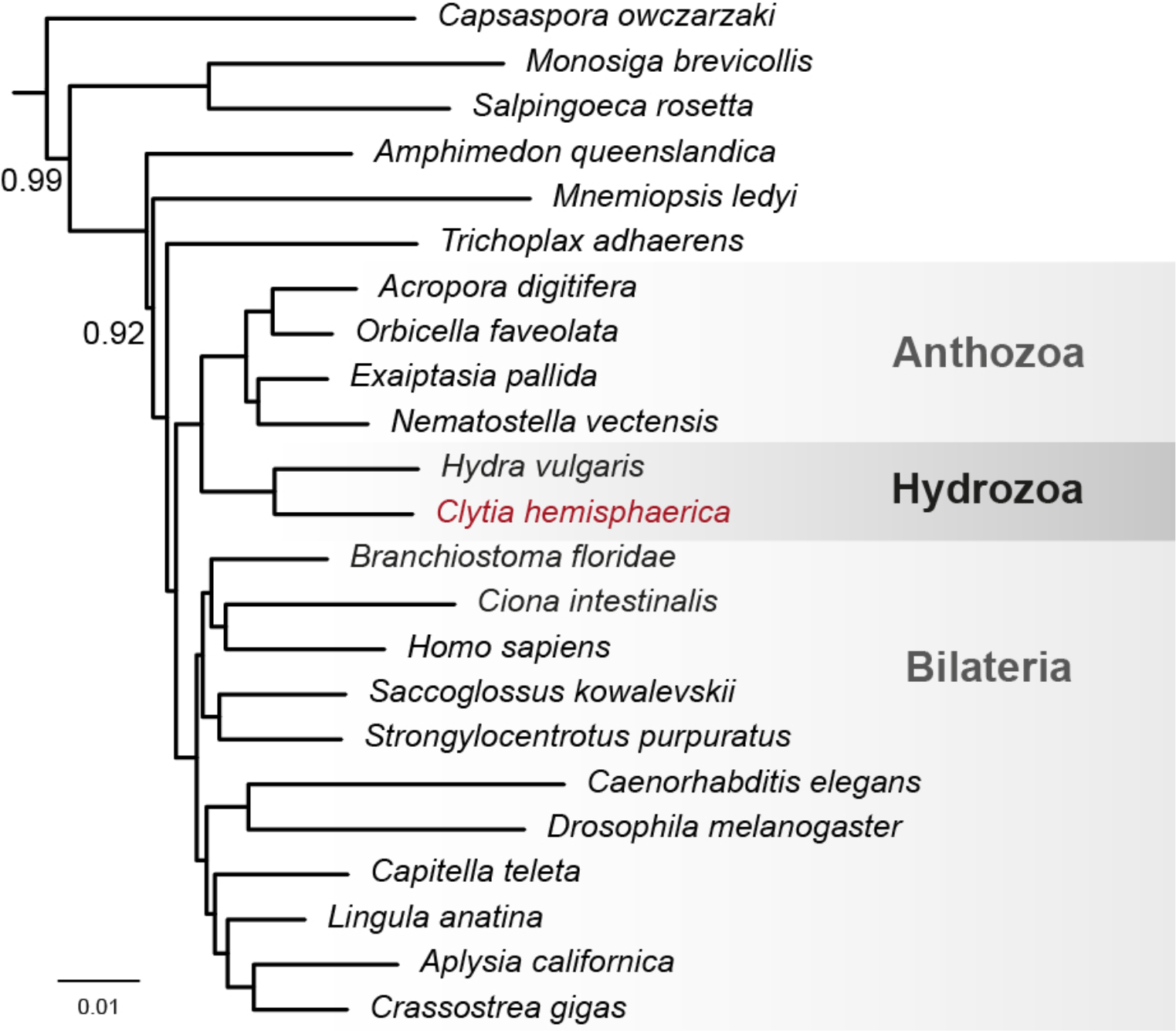
Phylogeny inferred using presence and absence of orthologous genes. The tree was rooted with *Saccharomyces cerevisiae* (not shown). All nodes have a posterior probability of 1 unless indicated.

Among many examples of gene gain in *Clytia*, we could identify new multigene families and also instances of Horizontal Gene Transfers (HGT), as illustrated by a UDP Glucose Dehydrogenase (UGDH) gene (fig. S3). UGDH is required for the biosynthesis of various proteoglycans and so to regulate signalling pathways during metazoan embryonic development [17]. Unusually, the *Clytia* genome contains two versions of the UGDH gene, including one acquired in Hydrozoa by HGT from a giant virus of the Mimiviridae family and expressed specifically during medusa formation. Interestingly, this UGDH xenolog was lost in the *Hydra* lineage and replaced by another UGDH acquired through HGT from bacteria (fig. S3) [10]. We also detected numerous gene duplications in the hydrozoan lineage, illustrated by the 39 Innexin gap junction genes (fig. S4), 14 Green Fluorescent Protein (GFP) and 18 Clytin photoprotein genes (fig. S5) found in the *Clytia* genome. The 4 GFPs and 3 Clytins previously reported in *Clytia* are transcribed from several recently duplicated genes, likely facilitating the levels of protein production needed to achieve the high cytoplasmic concentrations required for energy transfer between Clytins and GFPs [18].

Numerous likely gene losses in the hydrozoan lineage (i.e. absent in *Clytia* and *Hydra*) were confirmed by alignment-based phylogenetic analyses. These include many transcription factors involved in nervous system development in Bilateria (e.g. GBX, MNX, RAX, DBX, Pax3-7/PaxD, FOXG) [19,20]. Also absent were regulators of the anthozoan directive axis (the axis orthogonal to the oral/aboral axis, and likely related to the bilaterian dorsal/ventral axis [21,22], including HOX2 (represented in *Nematostella* by Anthox7/HoxC, Anthox8/HoxD), GBX, Netrin, the netrin receptor UNC-5, and chordin (The ‘Chordin-like’ gene described in *Hydra [23]*, is not orthologous to bilaterian and *Nematostella* Chordin [24]). Comparisons between the two available hydrozoan genomes revealed that 7 *Clytia* transcription factors that we identify as specifially expressed in the medusa (DRGX, PDX/XLOX, TLX, CDX, Six1/2, FoxL2, CnoxA, see below) have been lost in *Hydra*, but are still present in the transcriptome of one of its closest relatives possessing a medusoid stage, *Ectopleura larynx* (fig. S6). These *Hydra* gene losses thus likely relate to the loss of the medusa stage.

### Gene order disruption in the hydrozoan lineage

We tested conservation of gene order between *Clytia*, *Hydra*, *Nematostella* and *Branchiostoma floridae*, a bilaterian showing a particularly slow rate of loss of syntenic blocks [25], by identifying conserved adjacent pairs of orthologs (see methods) shared between two genomes. C*lytia* shares most genes in adjacent pairs with *Hydra* (340), including myc2 and its target CAD [26]. Fewer pairs were conserved between *Clytia* and either *Nematostella* (36) or *Branchiostoma* (16). Although *Nematostella, Hydra* and *Clytia*, as cnidarians, are equally distant phylogenetically from *Branchiostoma*, the number of genes in adjacent pairs in *Clytia*/*Branchiostoma* (16) or *Hydra*/ *Branchiostoma* (13) is considerably smaller than in *Nematostella*/*Branchiostoma* (110). Similar trends emerged from analyses limited to orthologs identified in all four species (Ch/Hv 51; Ch/Nv 8; Ch/Bf 4; Nv/Bf 20), so our conclusions are not biased by an inability to detect more divergent orthologs. These numbers are all significant compared to the same analyses performed with a randomized *Clytia* gene order. Such conservation of adjacent gene pairs possibly relates to coordinated transcription, or enhancers being embedded in adjacent genes [27]. In contrast, none of the previously described clusters of homologous developmental genes conserved between *Nematostella* and bilaterians, involving Wnt, Fox, NK, ParaHox or Hox family members [22,28,29] could be detected in *Clytia* (tab. S2) reinforcing the idea of rapid evolution of the genome organization in the common branch of *Clytia* and *Hydra*.

### Elevated stage specific gene expression in medusae and polyps

Hydrozoan life cycles are characterised by abrupt morphological transitions: metamorphosis from the planula to polyp; and growth and budding of the compex medusa from gonozooid polyps. To better understand global trends in differential gene use across the life cycle we produced a comprehensive replicated transcriptome dataset from 11 samples: early gastrula embryos, planulae at 3 developmental stages, newly-metamorphosed primary polyps, 3 components of the polyp colony (the feeding gastrozooid polyps, budding gonozooids and stolon), freshly-budded baby medusae, and adult male and female medusae (Fig. 1A).

Principal component analysis (PCA) of the most variably expressed genes across these transcriptomes confirmed sample reproducibility and revealed clear clustering of the three distinct hydrozoan life cycle stages: 1) the gastrula and planula samples 2) the polyp and stolon samples and 3) the medusa samples (Fig. 3A). Transcriptomes from gonozooids, specialized polyp structures containing developing medusae, were intermediate between the polyp and medusa ones. Inter-sample distances based on all genes presented a similar picture to the PCA (Fig. 3B). The main *Clytia* life cycle phases thus have qualitatively distinct overall profiles of gene expression, with a distance based dendrogram showing the polyp and medusa transcriptomes closer to each other than either is to the planula stage.

**Figure 3.**
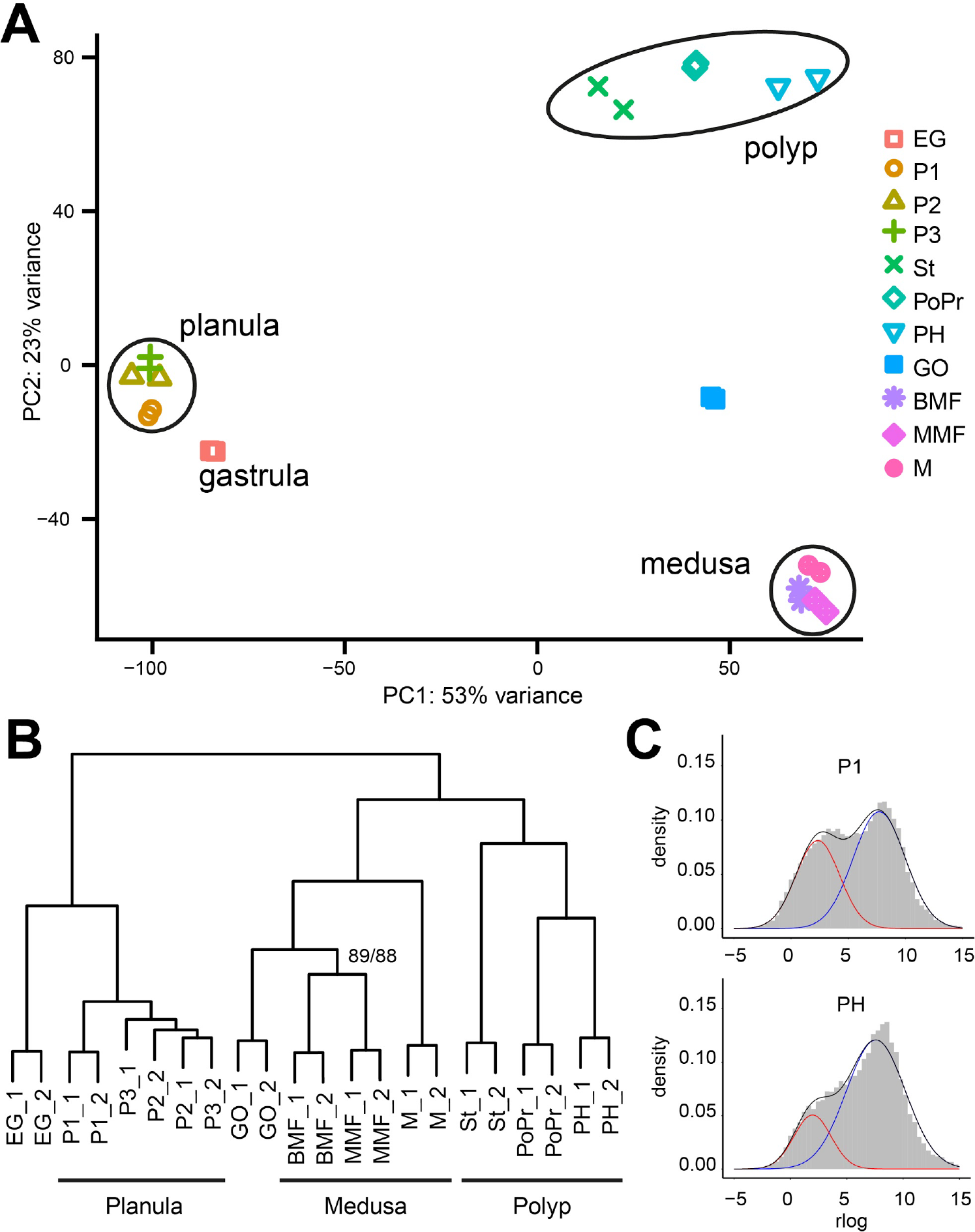
(A) PCA of all libraries from staged mRNA samples (B) Distance based clustering of all libraries. AU/BP values are 100 for all nodes unless indicated. Library names are as described in fig. 1 and methods. (C) Gene expression levels can be partitioned into on and off categories. Red and blue lines show fitted log-normal distributions and the black line their sum. Grey bars correspond to the empirically observed distribution of expression levels. Example libraries: P1, 1 day old planula; PH polyp head. All sample distributions are shown in fig. S7.

We used a novel method to identify stage specific genes, fitting the distribution of log-transformed gene expression for a given library to the sum of two Gaussian distributions, one of which we interpret as an ‘on’ set and the other an ‘off’ set (see methods, Fig. 3C, Fig. S7) [30]. By these criteria, polyp and medusa stages expressed more genes than embryo and planula stages, with highest expression of distinct genes at the primary polyp stage (19801 genes) and lowest expression at the early gastrula (13489 genes).

The majority of predicted genes, 84% (22472/26727) are expressed in at least one of our sampled stages (see methods; note that our gene prediction protocol includes data from deep sequencing of other mixed libraries), and 41% (10874/26727) are expressed at all stages. We defined genes as specific to a life-cycle stage if they were ‘on’ in at least one of the component samples for each main stage (Planula: early gastrula, 1, 2 or 3 day old planula; Polyp: stolon, primary polyp, or polyp head; Medusa: baby, mature or male), and ‘off’ in all other stages. We also identified genes specifically not expressed at either planula, polyp or medusa stages and so expressed at the two other stages. By these criteria, we found 335 planula, 1534 polyp and 808 medusa specific genes, and 1932, 284 and 981 genes specifically not expressed at planula, polyp and medusa stages respectively. We further filtered these data, requiring that genes also show significant (P<0.001, as defined by DESeq2 contrasts) expression differences between stages defined as ‘on’ and other stages, allowing a rigorous treatment of the variance between biological replicates. The application of this comparative expression criterion to the initial ″stage-specific″ gene sets derived by independent “on-off” determination for each sample, reduced the lists to 183 planula, 783 polyp and 614 medusa specific genes, and 1126, 180 and 374 genes specifically not expressed at planula polyp and medusa stages respectively. We can thus conclude that the two adult stages in the *Clytia* life cycle show greater complexity of gene expression than the planula larva.

In order to determine whether the medusa stage was enriched in genes found only in the medusozoan clade, as might plausibly be expected of an evolutionary novelty, we combined these lists of stage specific genes with a phylogenetic classification. We classified stage specific genes by the phylogenetic extent of the OMA Hierarchical Orthology Group of which they were a member (see methods), or the absence of significant sequence similarity to other species (for ‘*Clytia* specific’ sequences), and tested for enrichment of phylogenetic classes in each life-cycle stage (fig. S8). By these criteria, all three main life cycle stages (planula, polyp and medusa) were enriched in ‘*Clytia* specific’ sequences, indicating that phylogenetically ‘new’ genes are more likely than ‘old’ genes to show stage-specific expression, but are not associated with any one life cycle phase. In general, genes that evolved after the cnidarian/bilaterian split were more likely to be expressed specifically in adult (polyp/medusa) stages.

### Stage specific transcription factors

To address the nature of the molecular differences between stages, we assessed enrichment of Gene Ontology (GO) terms in stage specific genes relative to the genome as a whole. Planula larvae were found to be significantly enriched in G-protein coupled receptor signalling components, while polyp and medusa were enriched in cell-cell and cell-matrix adhesion class molecules (see tab. S3). Medusa specific genes were unique in being significantly enriched in the “nucleic acid binding transcription factor activity” term.

Confirming the strong qualitative distinction in gene expression profiles between planula, polyp and medusa (see above Fig. 3a and 3b) clustering of transcription factor expression profiles recovers the three major life-cycle stages (Fig. 4A). The majority of transcription factors (TFs) that were specific to a particular stage, using the criteria described above, were specific to the medusa (33, of which 11 are plausibly sex specific, see tab. S4). 12 were polyp specific (e.g. VSX, two HMX orthologs) and 16 were polyp-medusa specific, such that a total of 63 TFs were expressed at polyp and/or medusa stages but not at the planula stage (12.5% of the total TF). Only 3 TFs showed expression specific to the planula (TBX8 [31], a zinc finger containing protein and another T-box containing protein, neither obviously orthologous to other animal genes), while 8 were planula-polyp and 7 planula-medusa specific. This pattern is even more striking in the case of the 64 total homeodomain-containing TFs: 9 were medusa specific, 6 polyp specific and 5 polyp-medusa specific, with a total of 31% TF expressed at polyp and/or medusa stages but not at the planula stage. No homeodomain-containing TFs were identified as planula specific, with 1 specific to planula-polyp and 2 to planula-medusa.

**Figure 4.**
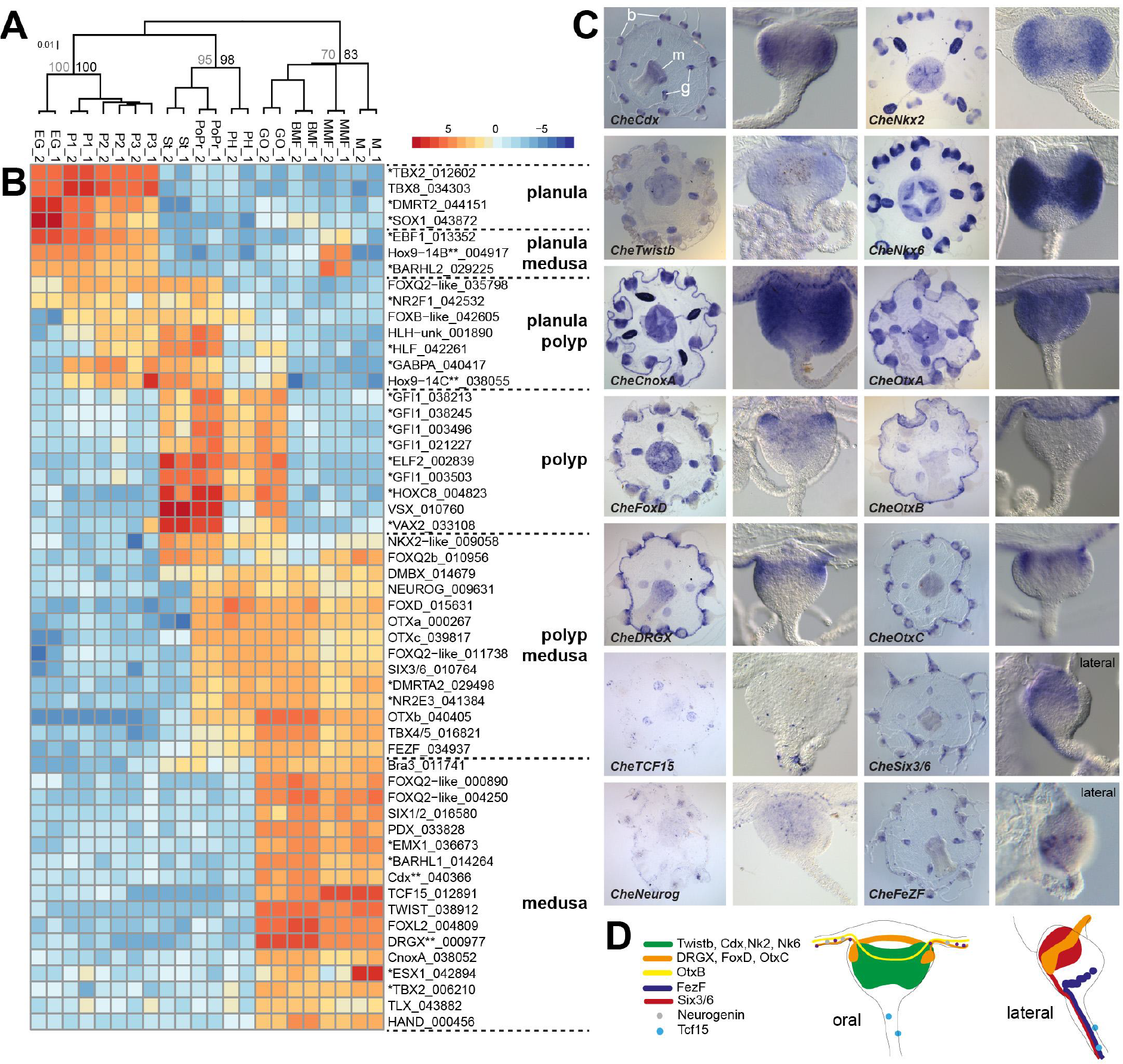
(A) Clustering of libraries by transcription factor expression distances. n=505 transcription factors. AU/BP values shown. (B) Heatmap showing major classes of transcription factor differential expression. Gene names are taken from orthology assignments of this work, with the suffix-like indicating assignment to a class but no precise orthology. In cases where assignment was not clear, names preceded by ^*^ are taken from best human Blast hits. Those followed by ^**^ are genes for which an assignment in *Clytia* existed in Genbank prior to this study (e.g. Cdx /Drgx). Trailing numbers are unique gene identifiers from this project. Expression units are rlog values from DESeq2 with the row means subtracted. Library names are as described in fig. 1 and methods. (C) *in situ* hybridization of medusa enriched TFs. (D) Oral and lateral view schematics of *Clytia* medusa tentacle bulbs indicating TF expression territories.

Among TFs expressed strongly in the medusa, but poorly at planula stages, we noted a large number with known involvement neural patterning during bilaterian development (Medusa only: Paraxis/TCF15, PDX/XLOX, CDX, TLX, Six1/2, DRGX, FOXQ2 paralogs; Polyp and Medusa: Six3/6, FoxD, FezF, OTX paralogs, HMX, TBX4/5, DMBX, NK2, NK6, Neurogenin1/2/3). We performed *in situ* hybridization analysis in medusae for a selection of these TF genes, and detected major sites of expression in the manubrium, gonads, nerve rings and tentacle bulbs (Fig. 4c,d). Within these structures, which are involved in mediating and coordinating feeding, spawning and swimming in response to environmental stimuli [1,32,33], expression showed a variety of distinct patterns, including scattered cell populations with putative sensory or effector roles. The variety of cellular distributions indicates a degree of complexity not previously described at the molecular level. We propose that, in *Clytia*, expression of conserved TFs in the medusa is associated with diverse cell types, notably with the neural and neurosensory functions of a complex nervous system, with continuous expression of certain transcription factors in post-mitotic neurons being necessary to maintain neuronal identity [34]. Members of the Sox, PRDL and Achaete scute (bHLH subfamily) orthology groups, commonly associated with neurogenesis [35,36] are detectable across all life cycle stages in *Clytia*, so our results are unlikely simply due to a higher production of nerve cells in the medusa.

Anthozoan larvae and bilaterian embryos express a common set of TFs at their respective aboral/ anterior ends, including Six3/6, FezF, FoxD, Otx, Rx, FoxQ2, and Irx [37,38] In the *Clytia* planula, whose anterior/aboral structures are relatively simple, most orthologs of this TF set are not expressed (Six3/6, FezF, FoxD, Otx orthologs), while another, Rx, was not found in the genome. A FoxQ2 gene (CheFoxQ2a) was reported to be aborally expressed in *Clytia* planulae [39] but this is a parlog of *Nematostella* aboral and *Platynereis* apical FoxQ2 [37,38], which are instead orthologous to CheFoxQ2b, a *Clytia* polyp-medusa specific gene (fig. S6.2, [39]). Irx is the only member of this conserved set of anterior/aboral TFs likely to be aborally expressed in *Clytia* planulae [40].

Organisation of the endoderm and oral ectoderm is also markedly simpler in *Clytia* than in *Nematostella* planulae, lacking mesenteries, mouth and pharyngeal structures. Correspondingly, many endoderm and mesoderm patterning genes expressed in many bilaterian larvae and *Nematostella* planulae (Cdx, PDX/XLOX, Nk2, Nk6, Twist, Paraxis/TCF15, Six1/2, Hand) [41,42], are not expressed in *Clytia* planulae. In contrast, despite different gastrulation mechanisms in anthozoans and hydrozoans, orthologs of TFs associated with gastrulation and endoderm formation in *Nematostella* [43], including FoxA, FoxB, Brachyury, Snail, Gsc are also expressed in oral-derived cells at gastrula and planula stages in *Clytia [40]*, as well as at polyp and medusa stage.

## Discussion

Three lines of evidence suggest that the *Clytia* genome has undergone a period of rapid evolution since the divergence of Hydrozoa from their common ancestor with Anthozoa (fig. 5). Firstly, rates of amino acid substitution appear to be elevated in hydrozoan relative to anthozoan cnidarians [44]. Secondly, orthologous gene content analysis shows that hydrozoans have the longest branches within the Cnidaria, with elevated rates of gene gain and loss (fig. 2). Thirdly, analysis of adjacent gene pairs shows more conservation between Anthozoa and *Branchiostoma* (as a representative of the Bilateria) than between Hydrozoa and *Branchiostoma*.

**Figure 5.**
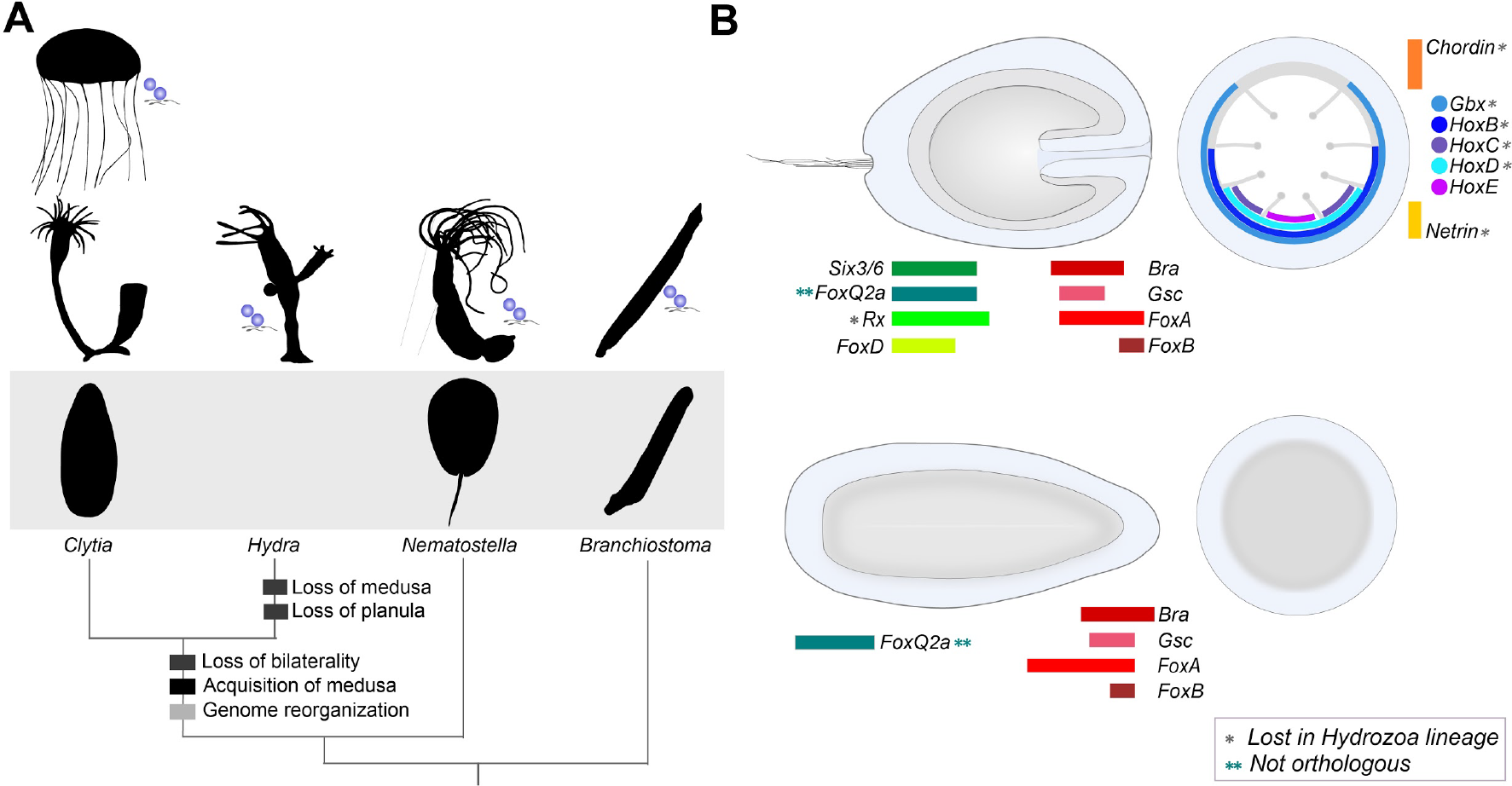
Simplification of polyp and planula stages in the hydrozoan lineage. (A) *Clytia hemisphaerica* displays the typical hydrozoan tri-phasic life cycle, comprising a planula stage, a (colonial) polyp stage, and a sexually reproducing medusa form. Both planula and medusa stages have been lost in the *Hydra* lineage. Hydrozoan planulae and polyps are morphologically simpler than those of Anthozoa (e.g. *Nematostella*), and have lost the bilateral symmetry. The comparison of *Clytia* and *Hydra* genomes with that of *Nematostella* shows that the hydrozoan lineage underwent important genome reorganization. (B) The planula larva of *Nematostella* (top) presents a well-defined endoderm and ectoderm, and bears an aboral apical organ. The eight internal mesenteries and the pharynx manifest the directive polarity axis, orthogonal to the oral-aboral one. A number of studies have identified a set of conserved transcription factors responsible for setting up the polarity axes and patterning the body. The *Clytia* planula (bottom) has a simpler morphology, with an ectodermal layer surrounding a mass of endodermal cells. Though the oral pole shares a set of developmental regulators with the planula of *Nematostella* (*Bra*, *Gsc*, *FoxA*, *FoxB*), the aboral pole appears to be highly divergent: expression of these known aboral transcription factors could not be detected – with the exception of *CheFoxQ2a* which does not belong to the *NvFoxQ2a* orthology group. Most of *Nematostella* directive axis regulators have been lost in Hydrozoa. Colored bars represent expression domains.

Gene expression analysis and lost developmental genes point to secondarily simplified planula and polyp structures in *Clytia*. The planula larva, in particular, shows an absence of key apical (i.e. aboral/anterior) and endomesoderm patterning genes considered ancestral on the basis of shared expression patterns in Anthozoa and bilaterian larvae [37,38,42]. Similarly, several genes with roles in patterning the directive axis of the anthozoan planula [21,45] are lost from the *Clytia* and *Hydra* genomes (Chordin, Hox2, Gbx, Netrin, unc-5), providing support for loss of bilaterality in medusozoans [24]. Much of the directive axis-patterning gene expression lost in *Clytia* planulae is, in *Nematostella*, likely involved in differentiating structures - mesenteries - that are maintained in the adult polyp, supporting the idea that the simple state of the *Clytia* polyp is secondary.

The medusa stage, as well as being morphologically complex, expresses a notable number of transcription factors that are conserved between cnidarians and bilaterians. These genes are expressed either specifically in the medusa (eg. DRGX, Twist and Pdx), or in both polyp and medusa but not planula stages (eg. Six3/6, Otx and FoxD), with medusa expression patterns suggesting roles in establishment or maintenance of neural cell-type identity. *Hydra* has lost the medusa from its life-cycle and has lost orthologs of most transcription factors that in *Clytia* are expressed specifically in the medusa, further supporting the notion that these genes are regulating the identity of cells now restricted to the medusa.

We propose then that, in part, the rapid molecular evolution we observe at the genome scale in Hydrozoa is connected as much to the simplified planula and polyp as the more obvious novelty of the medusa. Genomic and transcriptomic studies of additional medusozoan species, including scyphozoans, which produce medusae by strobilation and whose polyps are less simple than those of hydrozoans, may show whether the expansion of cell type and morphological complexity in the medusa phase, involving both old and new genes, has similarly been offset by reduction of gene usage in planula and polyp stages in other medusozoan lineages.

## Data availability

Data downloads and a genome browser are available at http://marimba.obs-vlfr.fr/

## Acknowledgments

We thank Iain Mathieson (UPenn) and Richard Mott (UCL) for statistical advice, and S. Collet, L Gissat and L. Gilletta for animal maintenance. Funding was provided by the CORBEL European Research Infrastructure cluster project, grants from the Agence Nationale de la Recherche (ANR-13-BSV2-0008-01 "OOCAMP" and ANR-13-PDOC-0016 "MEDUSEVO"), a Marie Curie training network (FP7-PEOPLE-2012-ITN 317172 "NEPTUNE"), André Picard Network, EMBRC-Fr as well as core CNRS and Sorbonne University funding to the LBDV.

## Methods

### Animals and extraction of genomic DNA

A 3-times self-crossed strain (Z4C)^2^ (male) was used for genomic DNA extraction, aiming to reduce polymorphisms. The first wild-type Z-strain colony was established using jellyfish sampled in the bay of Villefranche-Sur-Mer (France). Sex in *Clytia* is influenced by temperature [46] and some young polyp colonies can produce both male and female medusae. Male and female medusae from colony Z were crossed to make colony Z^2^. Two further rounds of self-crossing produced (Z4C)^2^ (see fig. S1 for relationships between colonies). For *in situ* hybridization (and other histological staining) we used a female colony Z4B, a male colony Z10 (offspring of (Z4C)^2^ x Z4B) as well as embryos produced by crossing Z10 and Z4B strains. (Z4C)^2^, Z4B and Z10 are maintained as vegetatively growing polyp colonies.

For genomic DNA extraction mature (Z4C)^2^ jellyfish were cultured in artificial sea water (RedSea Salt, 37‰ salinity) then in Millipore filtered artificial sea water containing penicillin and streptomycin for 3 to 4 days. They were starved for at least 24 hours. Medusae were snap frozen in liquid nitrogen, ground with mortar and pestle into powder then transferred into a 50 ml Falcon tubes (roughly 50~100 jellyfish/tube). About 20 ml of DNA extraction buffer (200 mM Tris-HCl pH 8.0 and 20 mM EDTA, 0.5 mg/ml proteinase K and 0.1% SDS) were added and incubated at 50°C for 3 hours until the solution became uniform and less viscous. An equal volume of phenol was added, vortexed for 1 minute, centrifuged for 30 minutes at 8000 g, then supernatant was transferred to a new tube. This extraction process was repeated using chloroform. X1/10 volume of 5M NaCl then 2.5 volumes of ethanol were added to the supernatant before centrifugation for 30 min at 8000 g. The DNA precipitate was rinsed with 70% ethanol, dried and dissolved into distilled water. 210 μg of DNA was obtained from 270 male medusae.

### Genome sequencing and assembly

Libraries for Illumina & 454 sequencing were prepared by standard methods (full details in supplementary methods file 1).

Sequence files were error corrected using Musket [47] and assembled using SOAPdenovo2 [48] with a large kmer size of 91 in an effort to separate haplotypes at this stage. We subsequently used Haplomerger2 to collapse haplotypes to a single more contiguous assembly [49]. Further scaffolding was done with L_RNA_Scaffolder using Trinity predicted transcript sequences (see below) [50,51].

### RNA extraction, transcriptome sequencing and gene prediction

RNA samples were prepared from Z4B female and Z10 male medusae and polyps, as well as embryos generated by crossing these medusae. Animals were starved for at least 24 hours before extraction and kept in Millipore filtered artificial sea water containing penicillin and streptomycin. Then they were put in the lysis buffer (Ambion, RNAqueous^®^ MicroKit), vortexed, immediately frozen in liquid nitrogen, and stored at -80°C until RNA preparation.

Total RNA was prepared from each sample using the RNAqueous^®^ Microkit or RNAqueous^®^ (Ambion). Treatment with DNase I (Q1 DNAse, Promega) for 20 min at 37°C (2 units per sample) was followed by purification using the RNeasy minElute Cleanup kit (Qiagen). See tab. S5 for total RNA (evaluated using Nanodrop). RNA quality of all samples was checked using the Agilent 2100 Bioanalyzer. The samples used to generate the expression data presented in fig. 3A are described in tab. S5. For the ‘mix’ sample, purification of mRNA and construction of a non-directional cDNA library were performed by GATC Biotech®, and sequencing was performed on a HiSeq 2500 sequencing system (paired-end 100 cycles). For the other samples, purification of mRNA and construction of a non-directional cDNA library were performed by USC Genomics Center using the Kapa RNA library prep kit, and sequencing was performed using either HiSeq 2500 (single read 50 cycles) or NextSeq (single read 75 cycles).

### Transcriptome assembly

We constructed a de novo transcriptome assembly without recourse to genomic data, assembling all RNA-Seq libraries with Trinity [50].

### Gene prediction

Genes were predicted from transcriptome data. Using tophat2, we mapped single end RNA-seq reads from libraries of early gastrula; 1, 2 and 3 day old planula; stolon, polyp head, gonozooid, baby medusa, mature medusa, male medusa, growing oocyte and fully grown oocyte to the genomic sequence [52]. In addition we mapped a mixed library made from the above samples but sequenced with 100bp paired end reads, and a further mature medusa library (100bp paired end). Genes were then predicted from these mappings using cufflinks and cuffmerge. Proteins were predicted from these structures using Transdecoder, and the protein encoded with the most exons taken as a representative for gene level analyses.

### Protein data sets

#### Protein analyses

We constructed a database of metazoan protein coding genes from complete genomes, including the major bilaterian phyla, all non-bilaterian animal phyla (including 6 cnidarian species) and unicellular eukaryotic outgroups. For the majority of species, we used annotation from NCBI, and selected one representative protein per gene, to facilitate subsequent analyses (tab. S6). We used the proteins as the basis for an OMA analysis to identify orthologous groups [53]. We converted the OMA gene OrthologousMatrix.txt file into Nexus format with datatype=restriction and used it as the basis for a MrBayes analysis, using corrections for genes present in fewer than 2 taxa ‘lset coding=noabsencesites|nosingletonpresence’, as described in [14]. This tree was then used in a subsequent OMA run to produce Hierarchical Orthologous Groups (HOGs). These HOGs were used as the basis for the phylogenetic classification of *Clytia* genes into one category out of eukaryotic, holozoan, metazoan, planulozoan, cnidarian or hydrozoan, based on the broadest possible ranking of the constituent proteins. Genes were presumed to have evolved in the most recent common ancestor of extant leaves, and leaves under this node where the gene was not present were presumed to be losses, with the minimum number of losses inferred to explain the observed presence and absence. *Clytia* specific genes were identified as those whose encoded proteins had no phmmer hits to the set of proteins used in the OMA analysis.

Where specific genes are named in the text, orthology assignments were taken from classical phylogenetic analysis (or in a few cases pre-existing sequence database names). Signature domains (e.g. Homeobox, Forkhead, T-box, HLH) were searched against the protein set using Pfam HMM models and hmmsearch of the hmmer3 package, with the database supplied ‘gathering’ threshold cutoffs [54,55]. Sequence hits were extracted and aligned with MAFFT [56] and a phylogeny reconstructed using RAxML with the LG model of protein evolution and gamma correction [57].

Transcription factors were assigned via matches beneath the ‘gathering’ threshold to Pfam domains contained in the transcriptionfactor.org database [58], with the addition of MH1, COE1_DBD, BTD, LAG1-DNAbind and HMG_Box.

*Ectopleura larynx* proteins were predicted with Transdecoder, including Pfam hit retention, from Trinity assembled reads (SRA accessions SRR923510_1 and SRR923510_2) [50,59].

#### Synteny analyses

Genes were ordered on their scaffolds (using the GFF files described in tab. S6) based on the average of their start and end position, and for each gene, the adjacent genes recorded, ignoring order and orientation but respecting boundaries between scaffolds (i.e. terminal genes had only one neighbour). Between species comparisons were performed using the Orthologous Groups from OMA, to avoid ambiguity from 1:many and many:many genes. When both members of an adjacent pair in one species were orthologous to the members of an adjacent pair in the other species, two genes were recorded as being involved in a CAPO (Conserved Adjacent Pair of Orthologs). A consecutive run of adjacent pairs (i.e. a conserved run of 3 genes) would thus be two pairs, but count as 3 unique genes.

#### RNA-Seq analyses

RNA-seq reads were aligned to the genome using STAR [60]. Counts of reads per gene were obtained using HTSeq-count. Gene level counts were further analyzed using the DESeq2 R package [61]. An estimate of the mode of row geometric means (rather than the default median) was used to calculate size factors. PCA and heatmaps were generated using regularized logarithms of counts (with blind = F). Bootstrapped hierarchical clustering was performed with pvclust using the default parameters [62]. Significant differences in gene expression were calculated via pairwise contrasts of different ‘conditions’ (replicated libraries). Planula stages were defined as any of 1,2 or 3 day old planula; Polyp any of stolon, primary polyp, polyp head or gonozooid; medusa any of gonozooid, baby medusa, mature medusa or male medusa. In order to be considered ‘up’ in planula, polyp or medusa, a gene needed to be significantly up (lfc threshold = 0.0, alt hypothesis = ‘greater’) in at least one ‘condition’ of that stage relative to all ‘condition’ of different stages.

#### Stage specific expression

To identify genes whose expression is restricted to particular stages we developed a two component pipeline: Firstly we discriminated ‘on’ vs ‘off’ genes independently for each stage as defined by absolute expression values, and then we assessed statistically significant differences between stages (which may not be between ‘on’ and ‘off’). To estimate for each sample the relative proportions of genes in the on vs. off categories, following Hebenstreit and Teichmann [30] we fitted each of our length normalized rlog-transformed gene expression datasets, averaged over replicates, to a mixture of two Gaussian distributions using the mixmodel R package [63]. The total number of ‘on’ genes for a given stage is estimated by multiplying the mixing proportion (lambda) of the ‘on’ peak by the total number of genes. Individual genes were defined as ‘on’ if they had a posterior probability > 0.5 of coming from the more highly expressed distribution. This approach avoids choosing an arbitrary “FPKM” (Fragments per Kilobase per Million mapped reads) value as an indicator of expression. Our frequency-distribution based approach defines gene ‘on’ or ‘off’ states independently of the total numbers of distinct transcripts expressed in a given sample, unlike FPKM values which are a measure of concentration and so for similarly expressed genes will be relatively higher in samples with low complexity.

#### GO term enrichment

GO terms were assigned via sequence hits to the PANTHER database using the supplied ‘pantherScore2.0.pl’ program. Term enrichment was tested using the ‘Ontologizer’ software with a ‘Parent-Child-Union’ calculation (the default) and Bonferroni multiple testing correction [64].

## Supplementary notes

### Note 1. Arguments for an ancestral medusa

Historically, some authors considered the medusa form to be the ancestral state of the cnidarians, with the anthozoans having lost this stage. Under this scenario hydrozoans are the earliest branching cnidarians (i.e. the sister group of other cnidarian taxa) because of their more radial (and hence assumed simpler) morphology, and consequently medusozoans are paraphyletic (to use modern terminology). Anthozoans would then have evolved from within the medusozoan group, and so must have lost the medusa stage [65]. Aside from debatable assumptions, this scenario has no support from molecular phylogenies [2,3]. More recently, some selected gene expression patterns have been used to support the idea of homology between the mesoderm of bilaterians and the entocodon of developing hydrozoan medusae, a ‘third’ cell layer that forms between ectoderm and endoderm during the early stages of medusa bud development [66–68]. Again, such arguments are not easily reconciled with modern phylogenies.

---

**Supplementary figure 1.**
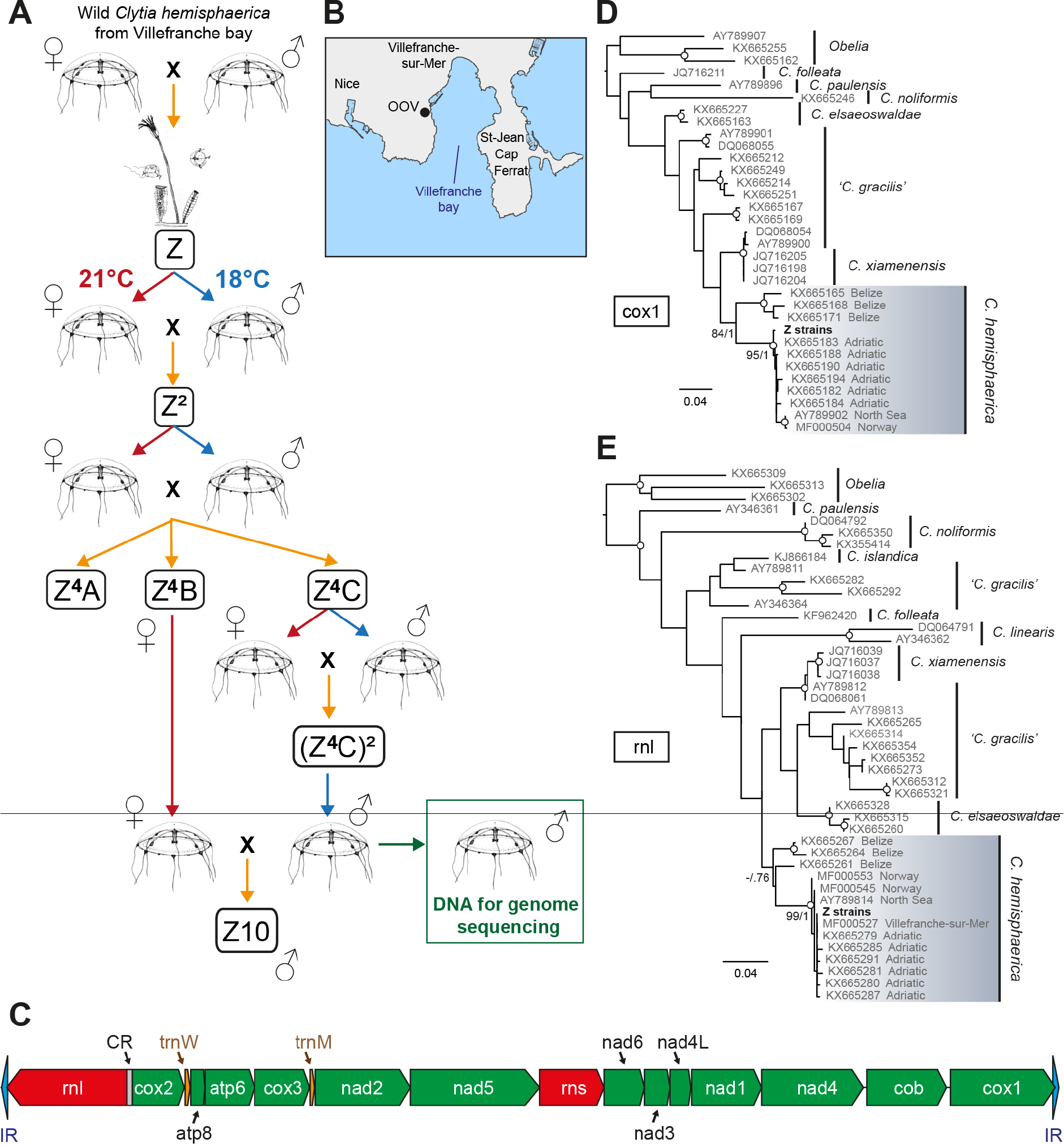
(A) Relationships between *Clytia hemisphaerica* laboratory strains; An original colony Z was derived by metamorphosis of a founder planula larva, with subsequent self crossing between male and female medusae budded from this colony. See methods for details. (B) The medusae used to generate the founder planula were collected in the bay of Villefranche-sur-Mer. (C) Organization of the linear mitochondrial genome of the sequenced Clytia Z strain showed the same organization as that inferred for the Hydroidolina common ancestor. (D,E) Maximum likelihood analyses using two mitochondrial phylogenetic makers (d: cox1/COI, e:rnl/16s RNA) unambiguously assign the Z strains to the species *Clytia hemisphaerica*.

**Supplementary figure 2.**
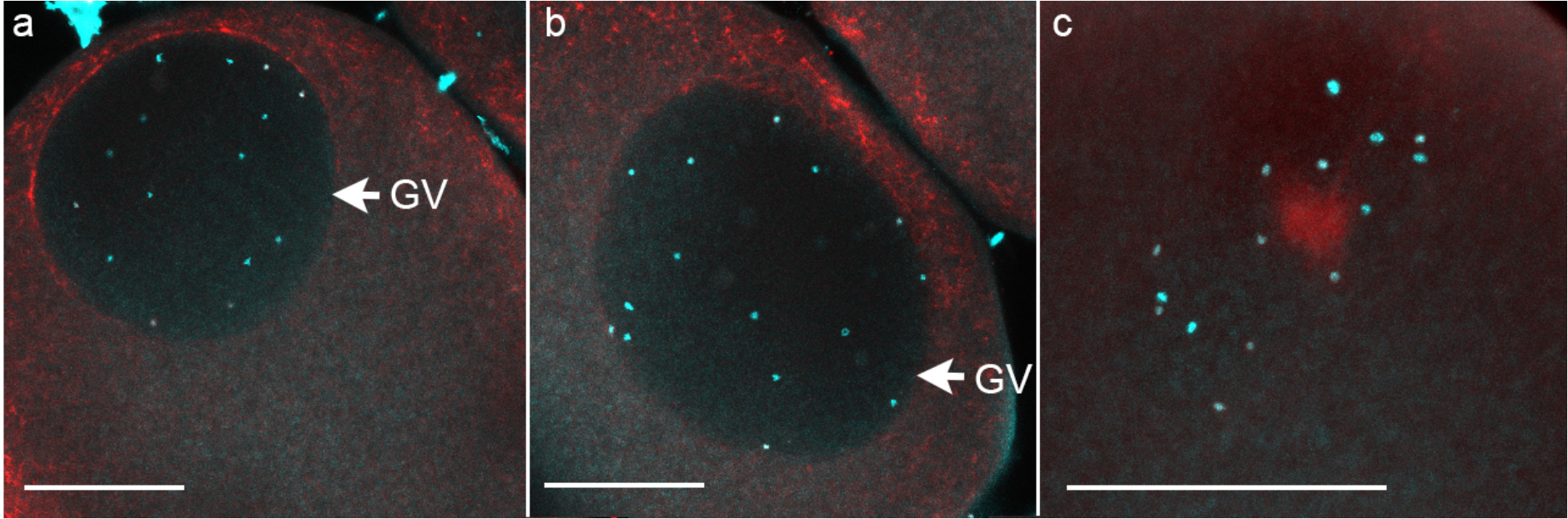
Hoechst staining (blue) of DNA in *Clytia* oocytes dissected from isolated gonads kept overnight in the dark and fixed, 35 minutes after exposing to light, for anti-tubulin immunofluorescence in red. The oocytes in a) and b), fixed using formaldehyde, are in the resting, meiotic prophase I arrested state, and t he duplicated chromosome pairs are clearly distinct in the dark oocyte nucleus (GV, Germinal Vesicle). The oocyte in c), fixed in cold methanol, has begun the maturation process, and the chromosomes are gathering on a microtubule aster. In each of 5 oocytes in which the Hoechst-stained lumps were clearly distinct, 15 chromosome packets were counted. Maximum projections of z stacks covering the whole GV or chromosome cluster are shown. Scale bar 50μm.

**Supplementary figure 3.**
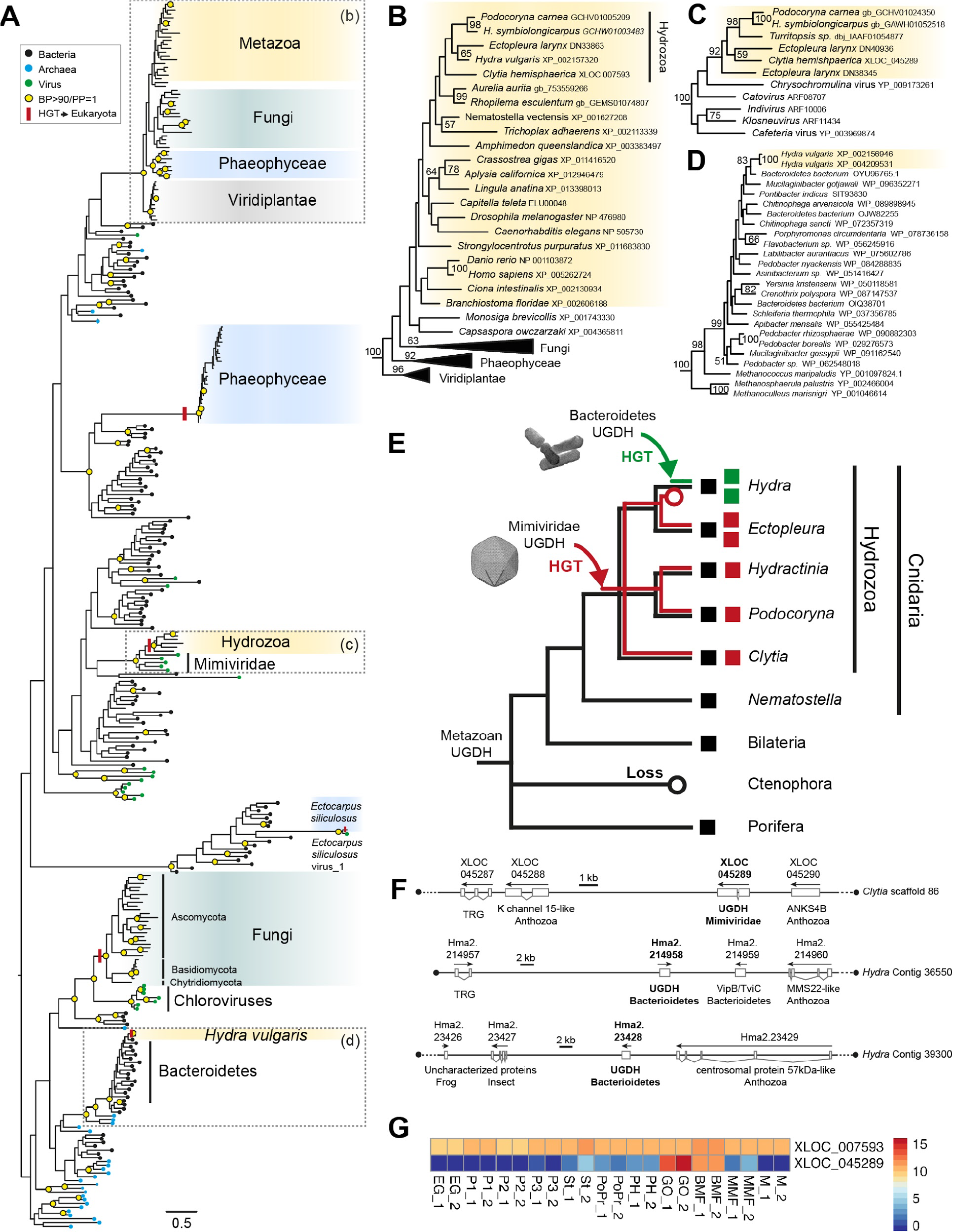
(A) Maximum likelihood phylogenetic reconstruction (RaxML, LG+Gamma) of the two UGDH-like (UDP-glucose 6-dehydrogenase) genes found in the *Clytia* genome. (B) Metazoa UGDH subtree including one *Clytia* ortholog. (C) Subtree showing that the second *Clytia* UGDH, together with several other hydrozoan sequences, are nested within UGDH genes from mimiviridae, strongly arguing for a Horizontal Gene Transfer (HGT), to a hydrozoan ancestor, from a giant virus of the Mimiviridae family. (D) Subtree showing that two UGDH-like genes of *Hydra* are nested within a strongly supported Bacteroidetes clade, strongly suggesting that they were acquired in the *Hydra* lineage by HGT from bacteria. (E) Evolutionary scenario for the evolution of the UGDHs found in the Hydrozoa lineage. (F) genomic location of *Clytia* and *Hydra* UGDH genes inherited by HGT. Presence of clear metazoan orthologs in these scaffolds argues against bacterial/viral contamination. Gene name and taxon of the BLASTp hit are indicated below the genes; TRG: Taxon Restricted Gene - no BLASTp hit. (G) expression level (rlog values) of the two *Clytia* UGDH genes across the life cycle (see legend of fig. 1 for details on stage abbreviations). Note the ubiquitous expression of the “metazoan” UGDH gene (XLOC_007593) and the Gonozoid (GO) and Baby Medusae (BMF) specific expression of the “Mimiviridae-related” UGDH (XLOC_045289).

**Supplementary figure 4.**
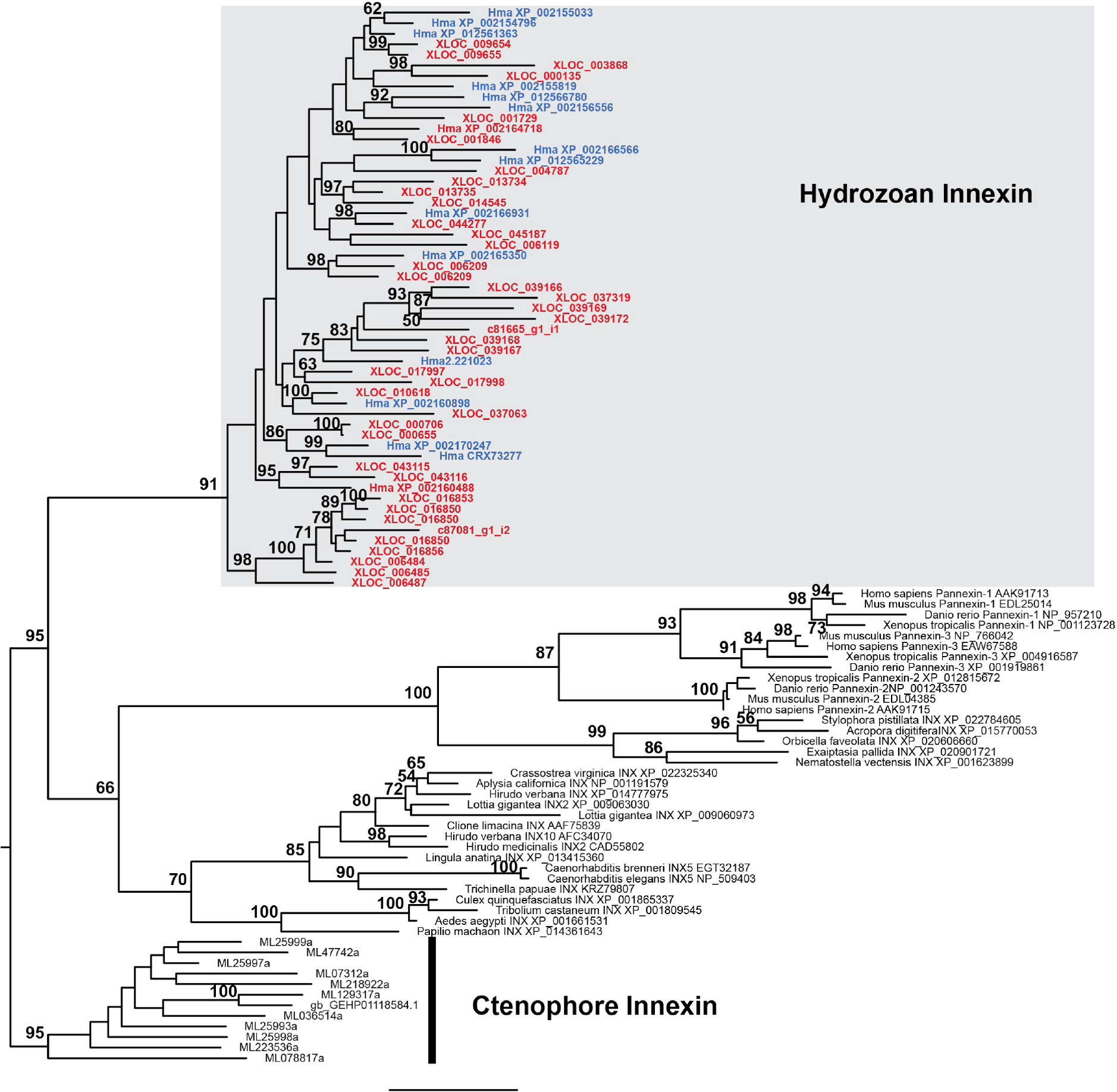
Innexin Maximum Likelihood phylogeny. Model LG+Gamma RAxML. Rooted with ctenophore innexin. Red: *Clytia* sequences, Blue: *Hydra* sequences. Values on nodes = rapid RAxML Bootstrap values (500 replicates) > 50%.

**Supplementary figure 5.**
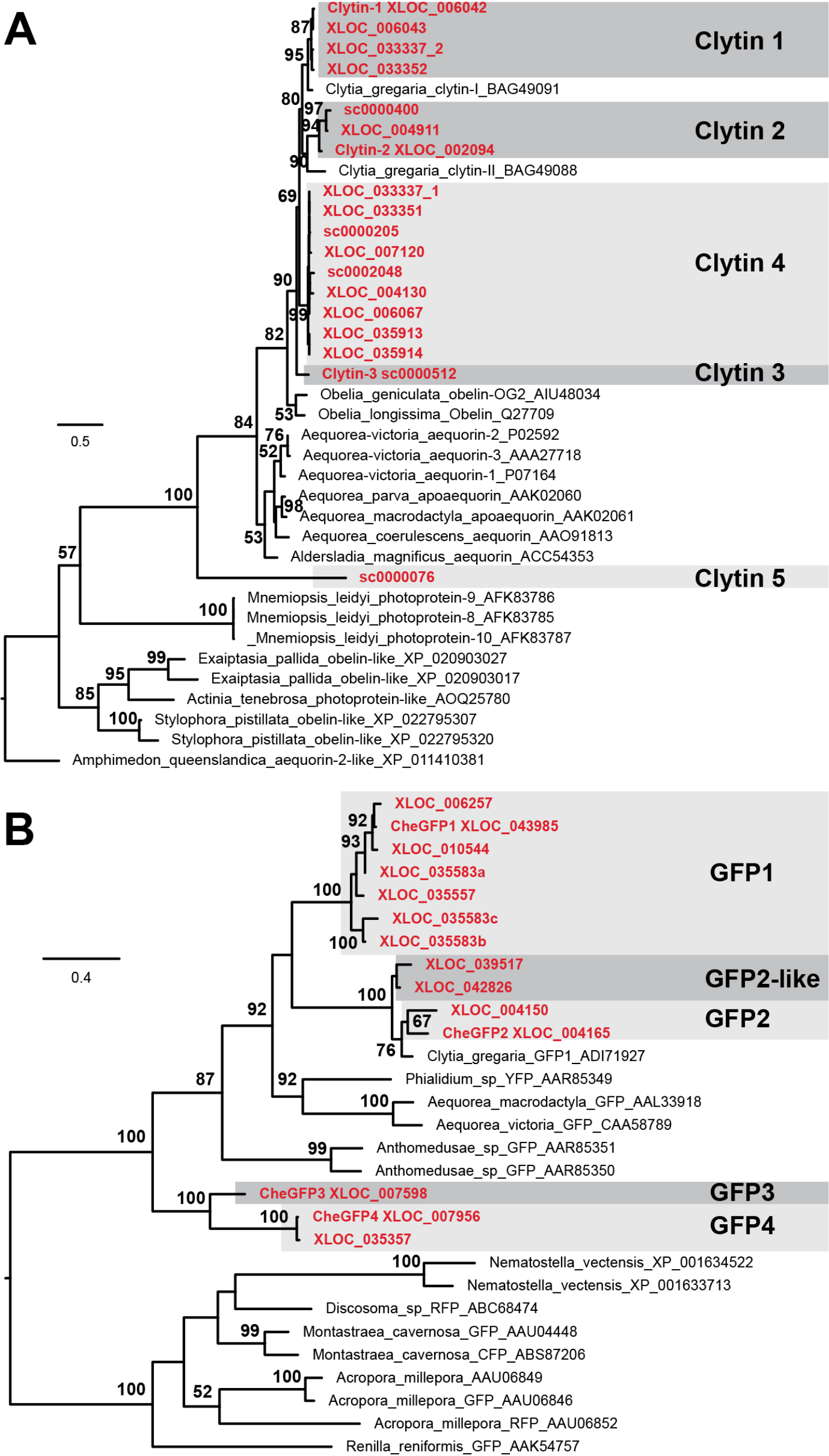
Maximum likelihood phylogenetic analysis of *Clytia* and Clytin-like (A) and GFP-like (B) genes. Model LG+Gamma RAxML. Rooted with sponge aequorin-like (A) and anthozoan GFP-like (B) genes. Values on nodes = rapid RAxML Bootstrap values (500 replicates). Only BP>50% are labeled. Red: Clytia sequences.

**Supplementary figure 6.1.**
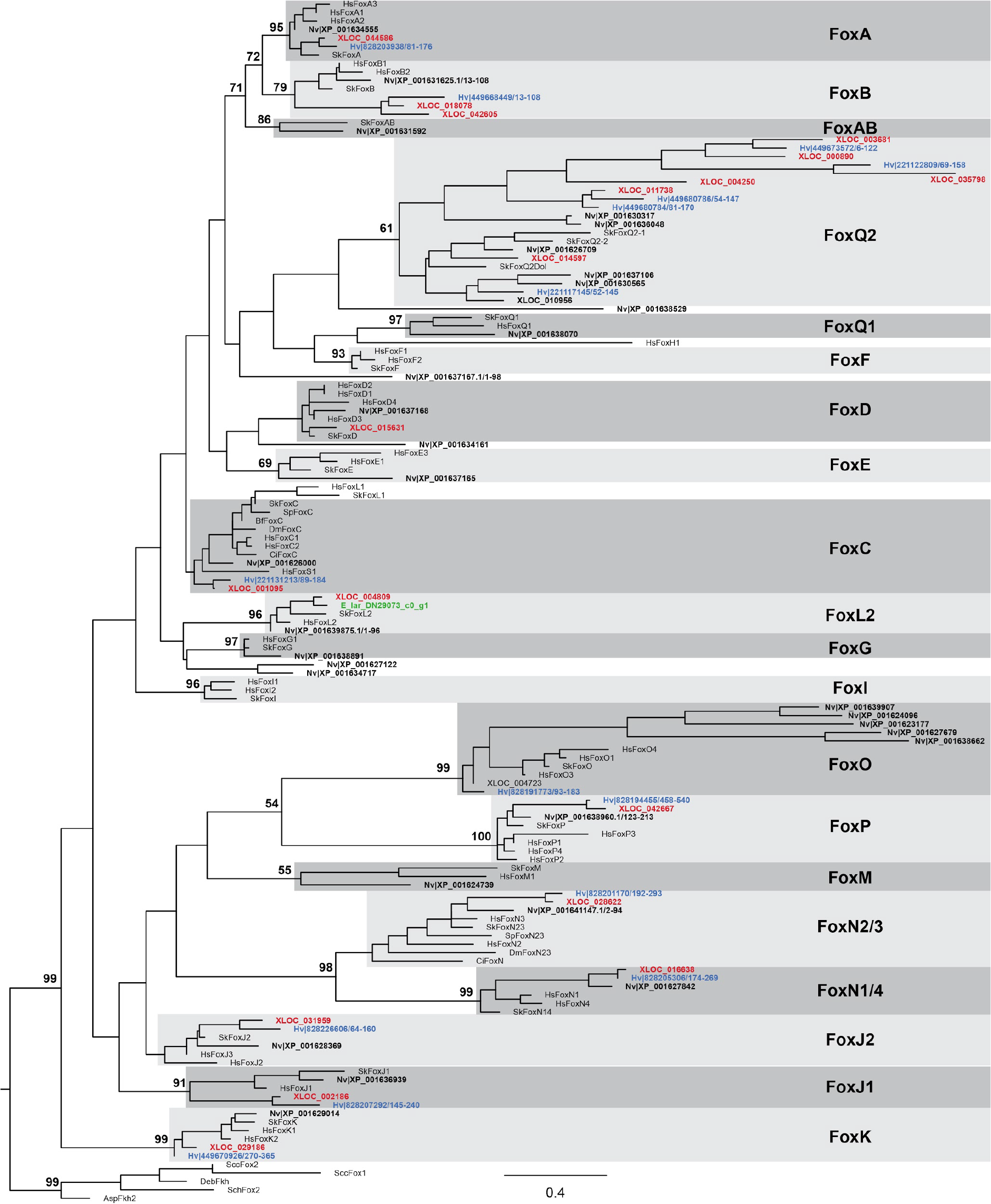
Maximum likelihood phylogenetic analysis of *Clytia* and *Hydra* Fox-like genes. Model LG+Gamma RAxML. Rooted with non metazoan Forkhead domains. Values on nodes = rapid RAxML Bootstrap values (500 replicates). Only BP>50% for nodes outside Fox families are labeled. From this topology it can be inferred that FoxM, FoxG, FoxQ1 were lost in the common ancestor of *Clytia* and *Hydra* and FoxL2, FoxD were lost in the *Hydra* lineage. Red: *Clytia hemisphaerica*, Green: *Ectopleura larynx* FoxL2, black bold: *Nematostella vectensis,* Blue: *Hydra* - Hm: *Hydra magnipapillata*, Hv: *Hydra vulgaris*, Sk: *Saccoglossus kowalevskii*, Hs: Homo sapiens.

**Supplementary figure 6.2.**
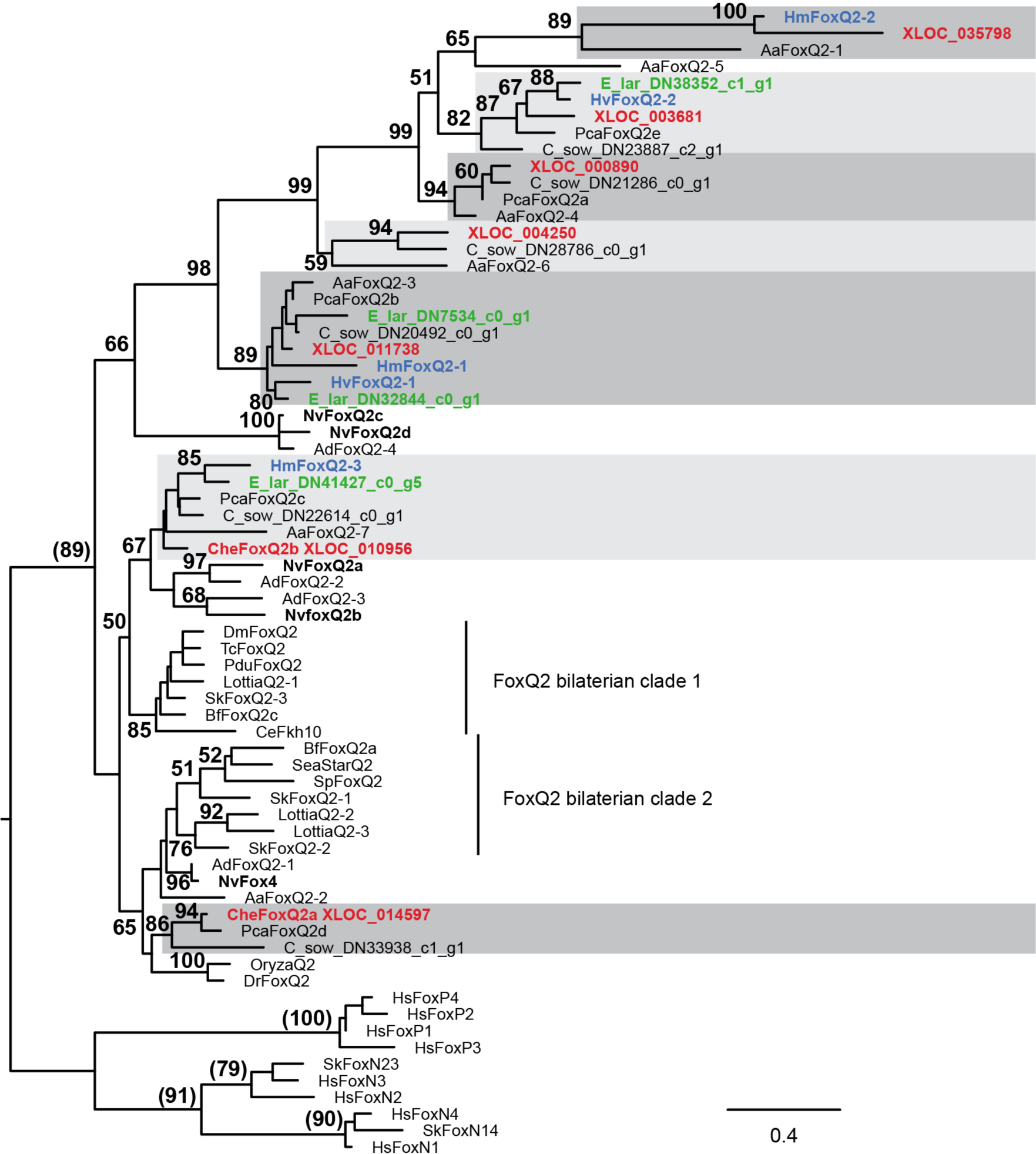
Maximum likelihood phylogenetic analysis of *Clytia* and *Hydra* FoxQ2-like genes. (Model LG+Gamma RAxML) rooted with FoxP/FoxN14/FoxN23. Rooted and unrooted ML analyses lead to the exact same ingroup topology. rapid RAxML Bootstrap values >50% from the ML analysis of the unrooted dataset are indicated next to the branches. BP values between brackets are from the rooted ML analysis. From this topology, it can be inferred that the cnidarian ancestor had at least 3 FoxQ2-like genes, two of each were also present in the bilaterian ancestor. Expansion of one of those 3 ancestral FoxQ2 gene in the common ancestor of Medusozoa lead to the formation of 7 FoxQ2 medusozoan orthologous groups (highlighted in grey), all still present in the *Clytia* genome. The *Hydra* lineage lost 3 of those FoxQ2 paralogs. Note that the aboraly expressed *Clytia* FoxQ2a does not belong to the same FoxQ2 orthologous group as the aboraly expressed *Nematostella* FoxQ2a. Red: *Clytia hemisphaerica* sequences, Green: *Ectopleura larynx* sequences, black bold: *Nematostella vectensis,* Blue: Hydra sequences, Hm: *Hydra magnipapillata*, Hv: *Hydra vulgaris*, Pca: *Podocoryna carnea*, C_sow: *Craspedacusta sowerbyi*, Aa: *Aurelia aurita*, Sk: *Saccoglossus kowalevskii*, Hs: Homo sapiens, Lottia: *Lottia gigantea*, Sp: *Strongylocentrotus purpuratus*, Bf: *Branchiostoma floridae*, Pdu: *Platynereis dumerilii*, Dm: *Drosophila melanogaster*, Tc: *Tribolium castaneum*, Ce: *Caenorhabditis elegans.*

**Supplementary figure 6.3.**
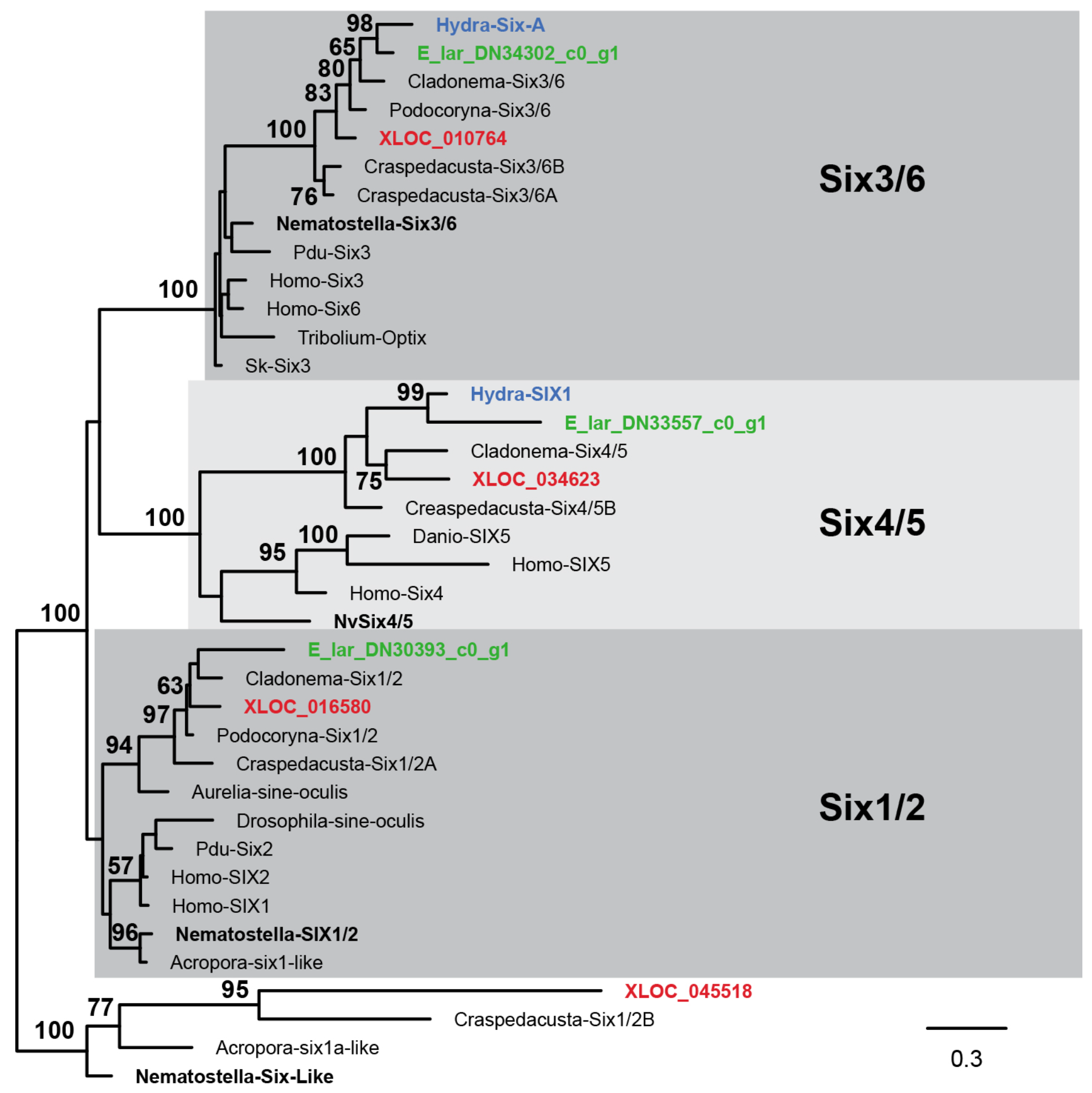
Maximum likelihood phylogenetic analysis of *Clytia* and *Hydra* Six genes. (Model LG+Gamma RAxML) rooted with distantly related Six-like genes found in cnidarians. Values on nodes = rapid RAxML Bootstrap values (500 replicates). From this topology it can be inferred that Six1/2 was lost in the *Hydra* lineage after its divergence with the *Ectopleura* lineage. Red: *Clytia hemisphaerica*, black bold: *Nematostella vectensis*, Blue: *Hydra magnipapillata*, Green: *Ectopleura larynx*. Pdu: *Platynereis dumerilii*, Sk: *Saccoglossus kowalevskii*.

**Supplementary figure 6.4.**
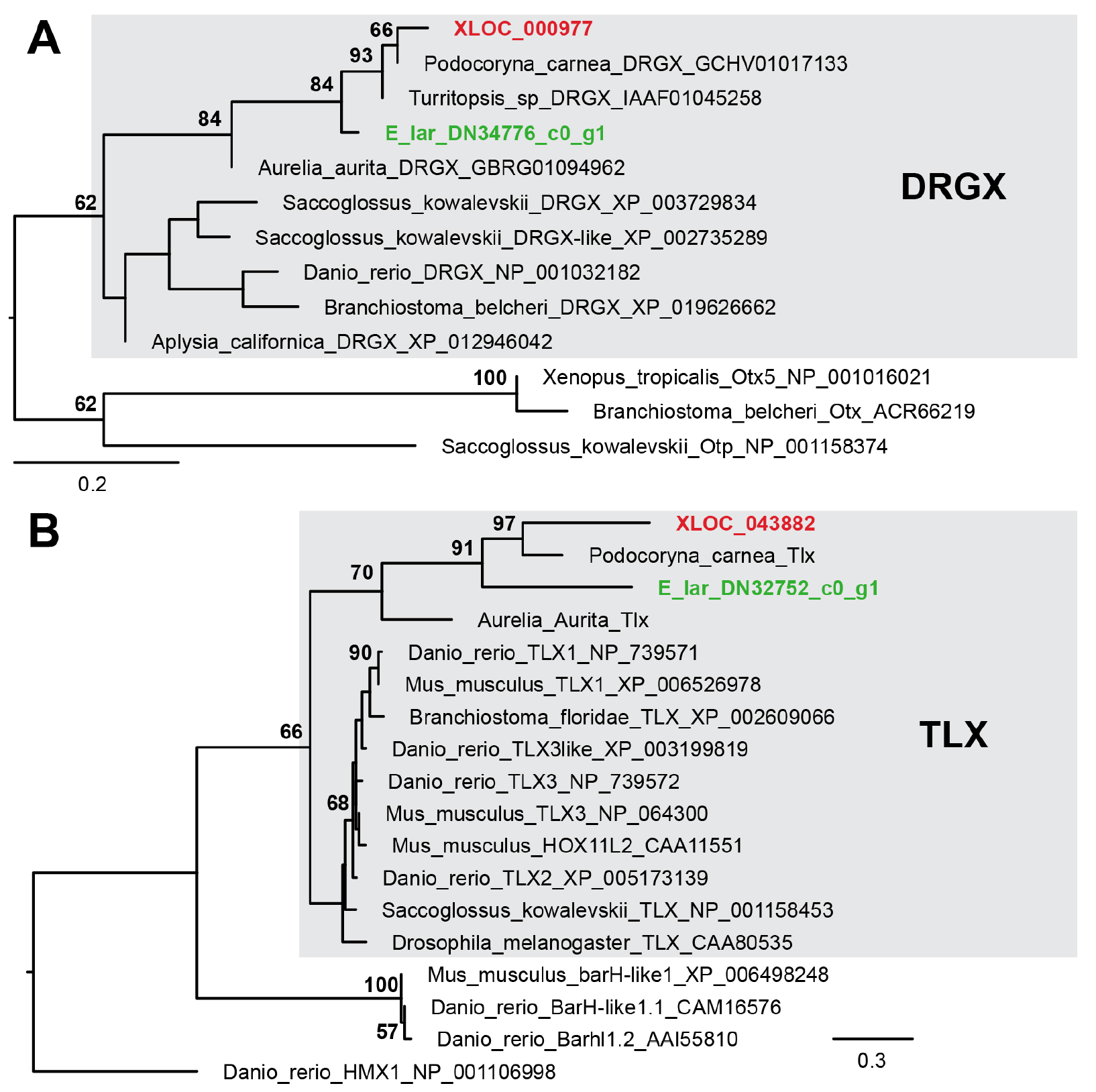
Maximum likelihood phylogenetic analysis of *Clytia and Ectopleura* and DRGX (A) and TLX (B) genes. Model LG+Gamma RAxML. Values on nodes = rapid RAxML Bootstrap values (500 replicates). Only BP>50% are labeled. Red: Clytia sequences.

**Supplementary figure 6.5.**
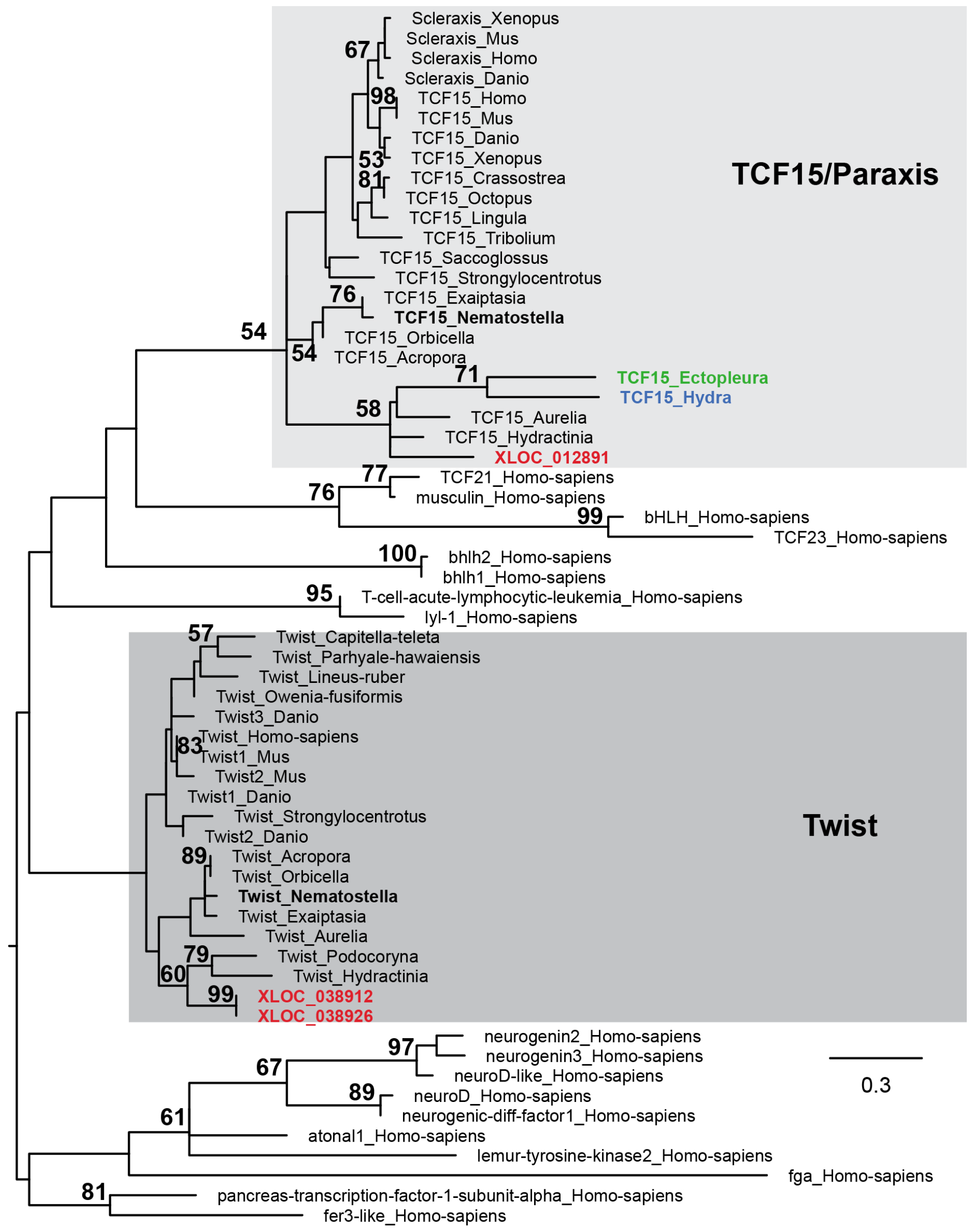
Maximum likelihood phylogenetic analysis of *Clytia* Twist and TCF15 genes. (Model LG+Gamma RAxML) rooted with a selection of human bHLH genes. Values on nodes = rapid RAxML Bootstrap values (500 replicates). From this topology it can be inferred that Twist was lost in a Hydra ancestor. Red: *Clytia hemisphaerica*, black bold: *Nematostella vectensis*, Blue: *Hydra magnipapillata*, Green: *Ectopleura larynx*.

**Supplementary figure 6.6.**
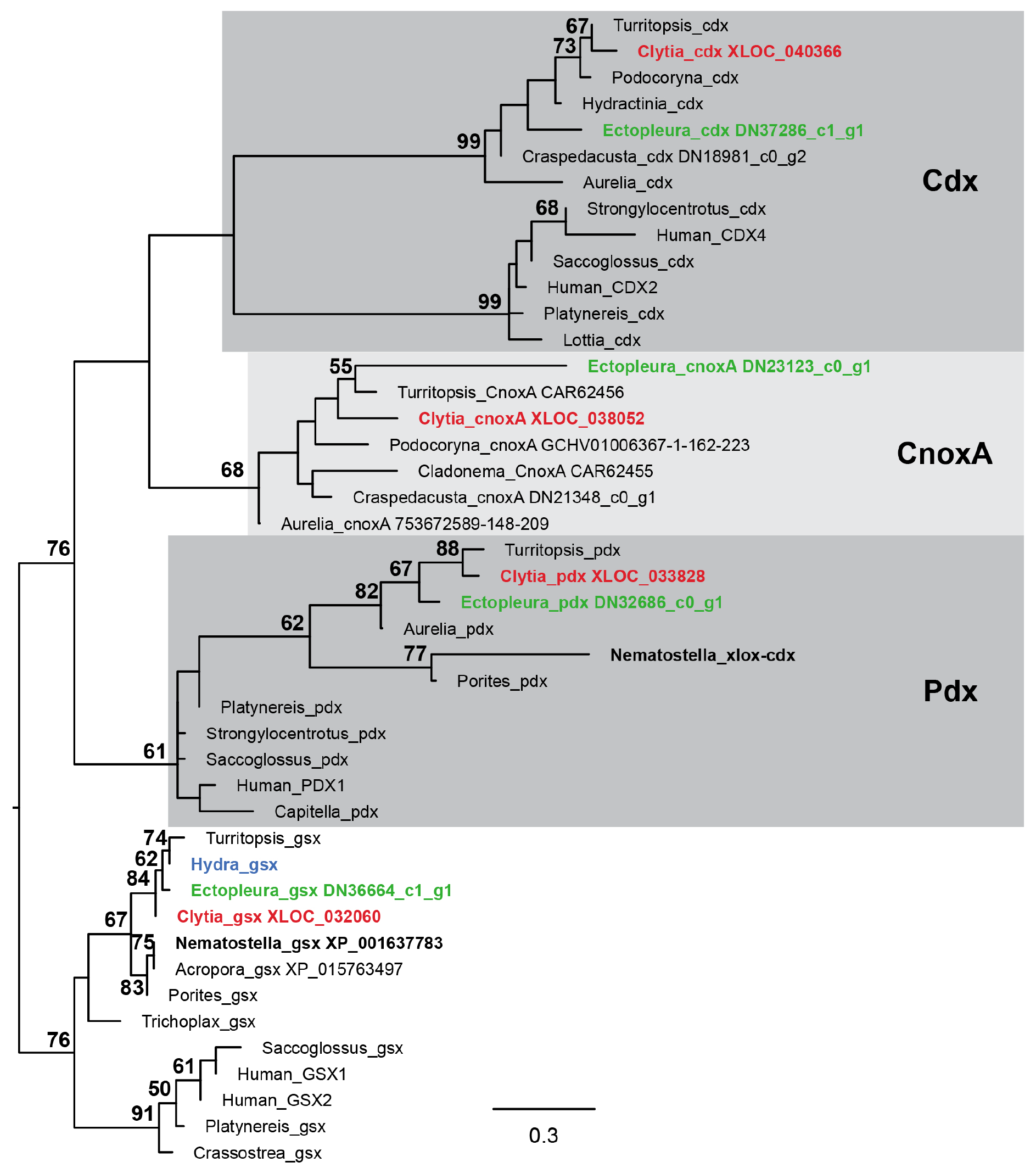
Maximum likelihood phylogenetic analysis of *Clytia* Cdx, Pdx, CnoxA and Gsx genes. (Model WAG+Gamma RAxML) rooted with Gsx genes. Values on nodes = rapid RAxML Bootstrap values (500 replicates). From this topology it can be inferred that Cdx, Pdx and CnoxA were lost in the *Hydra* lineage after the divergence with *Ectopleura*. Red: *Clytia hemisphaerica*, black bold: *Nematostella vectensis*, Blue: *Hydra magnipapillata*, Green: *Ectopleura larynx*.

**Supplementary figure 6.7.**
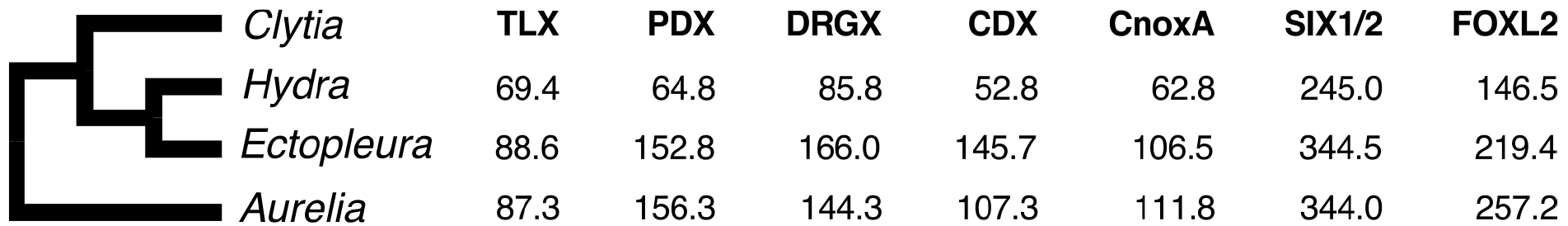
Evidence for gene loss in *Hydra*. Phylogenetic analysis shows orthologs of the named *Clytia* genes in *Ectopleura* and *Aurelia* but not in *Hydra*. The table gives the phmmer bit score [54] for the pairwise comparison of the best hit of the *Clytia* gene in these three medusozoan species. The *Hydra* best hits are non-orthologous and substantially lower scoring than the medusoid producing sister species (*Ectopleura larynx*), or more phylogenetically distant species that retain an ortholog and medusa stage (the scyphozoan *Aurelia aurita*).

**Supplementary figure 7.**
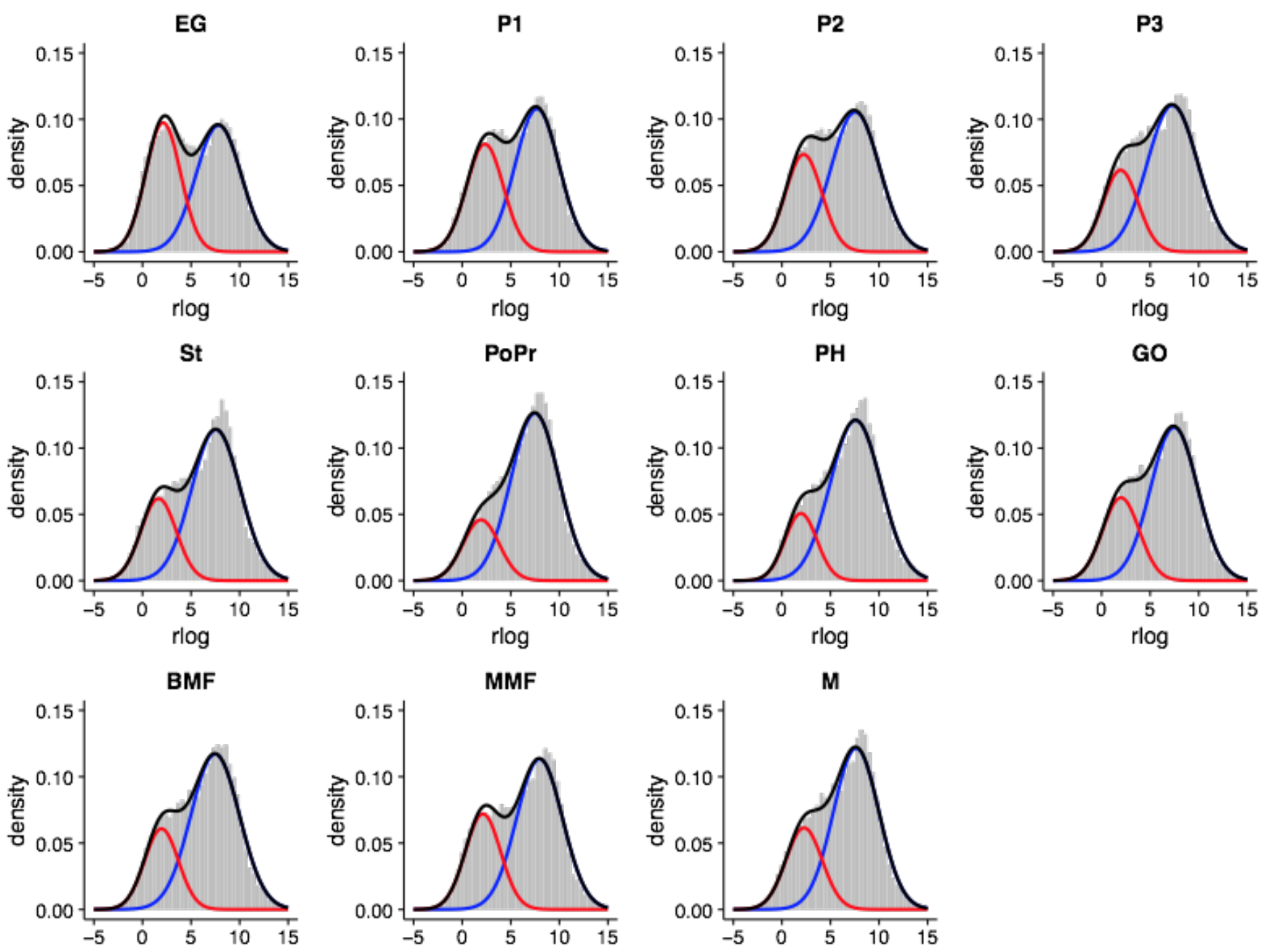
Gene expression for all life-cycle stages partitioned into ‘on’ and ‘off’ distributions. Red and blue lines show fitted log-normal distributions and the black line their sum. Grey bars correspond to the empirically observed distribution of expression levels. Library names are as for fig. 1a.

**Supplementary figure 8.**
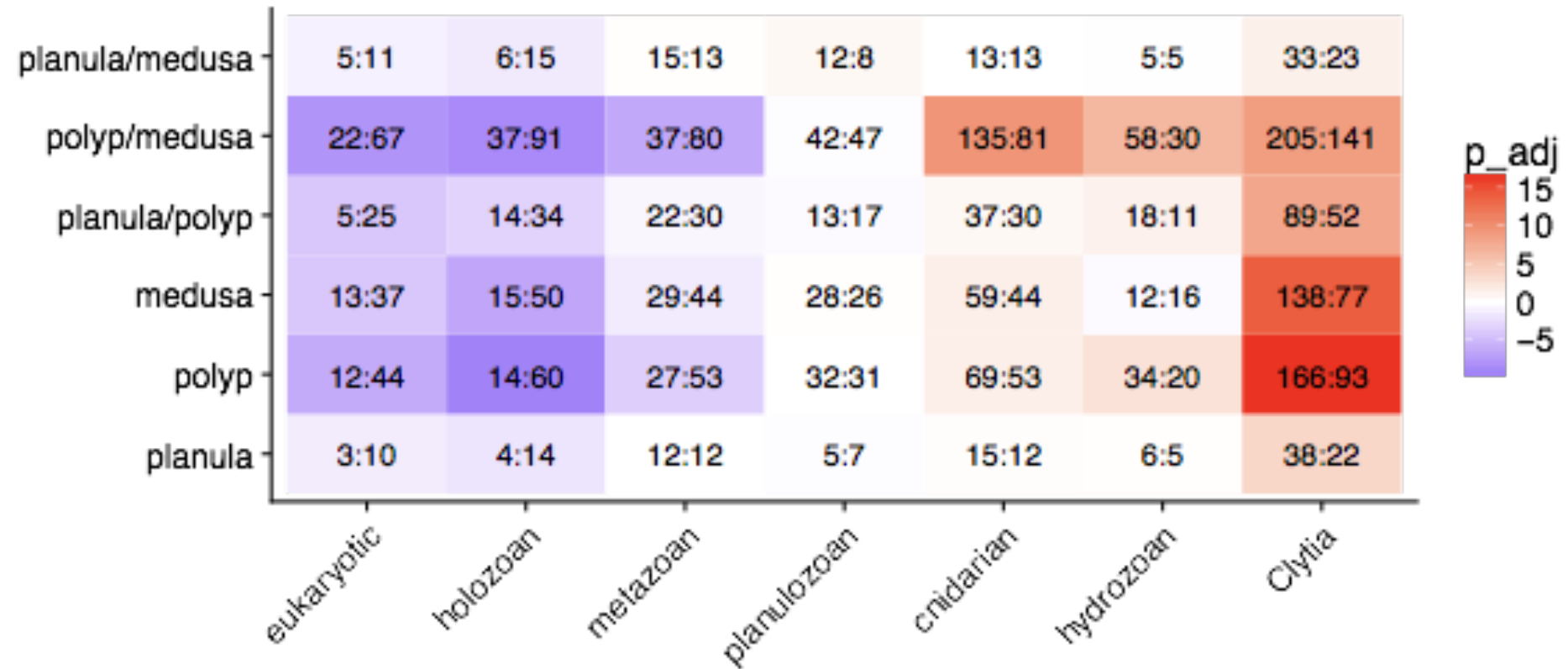
Enrichment of OMA HOG phylogenetic class across stage specific genes. Each cell contains the ratio of observed to expected genes for that stage specificity (vertical axis) and phylogenetic scope, as determined by OMA Hierarchical Orthologous Group composition (horizontal axis). Red cells show an enrichment of the phylogenetic class, blue cells a depletion. Effect significance was determined by a Chi-squared test for each cell, corrected for multiple testing using the Benjamini and Hochberg (FDR) method of the p.adjust R function. These values were used to determine the cell color after log_10_ rescaling, and under-represented cells being made negative.

**Supplementary Table 1.** Comparison between *Clytia*, *Hydra* and *Nematostella* genomic features

| Species | *Clytia hemisphaerica* | *Hydra vulgaris* | *Nematostella vectensis* |
| --- | --- | --- | --- |
| Clade | Hydrozoa | Hydrozoa | Anthozoa |
| Presence of jellyfish | yes | no | no |
| Chromosome number (n) | 15 | 15 | 15 |
| Genome size (Mb) | 445 | 1,050 | 457 |
| Scaffold N50 (kb) | 366 | 92.5 | 472 |
| GC content (%) | 35 | 29 | 39 |
| Repeats (%) | 41 | 57 | 26 |
| Number of genes | 26,203 | 31,452 | 27,273 |
| BUSCO (%) | 86.4 | 76.8 | 91.4 |

**Supplementary Table 2.** Expression patterns and genome locations for all transcription factors.

**Supplementary Table 3.**
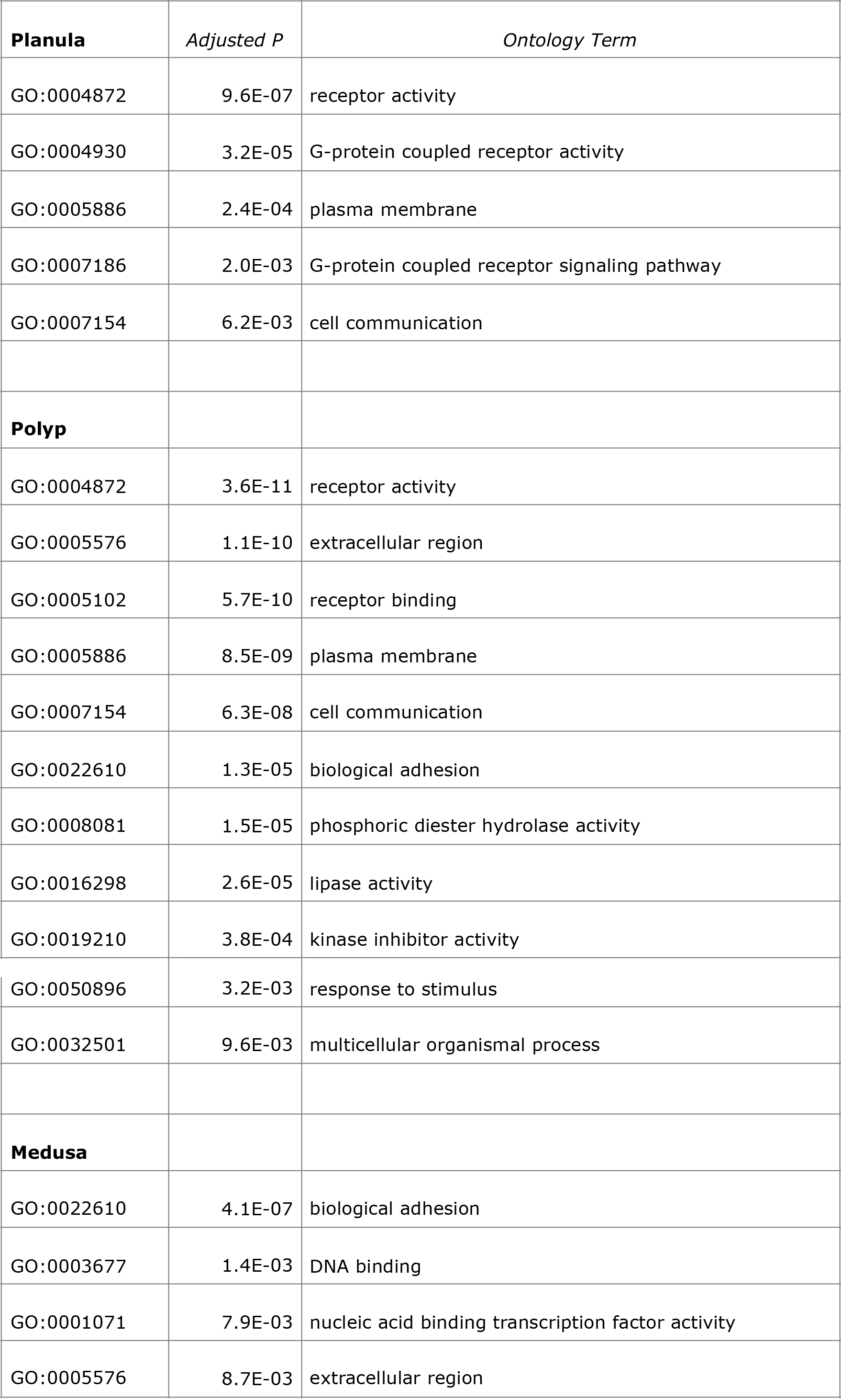
Significant gene ontology enrichments at major life-cycle stages.

**Supplementary Table 4.**
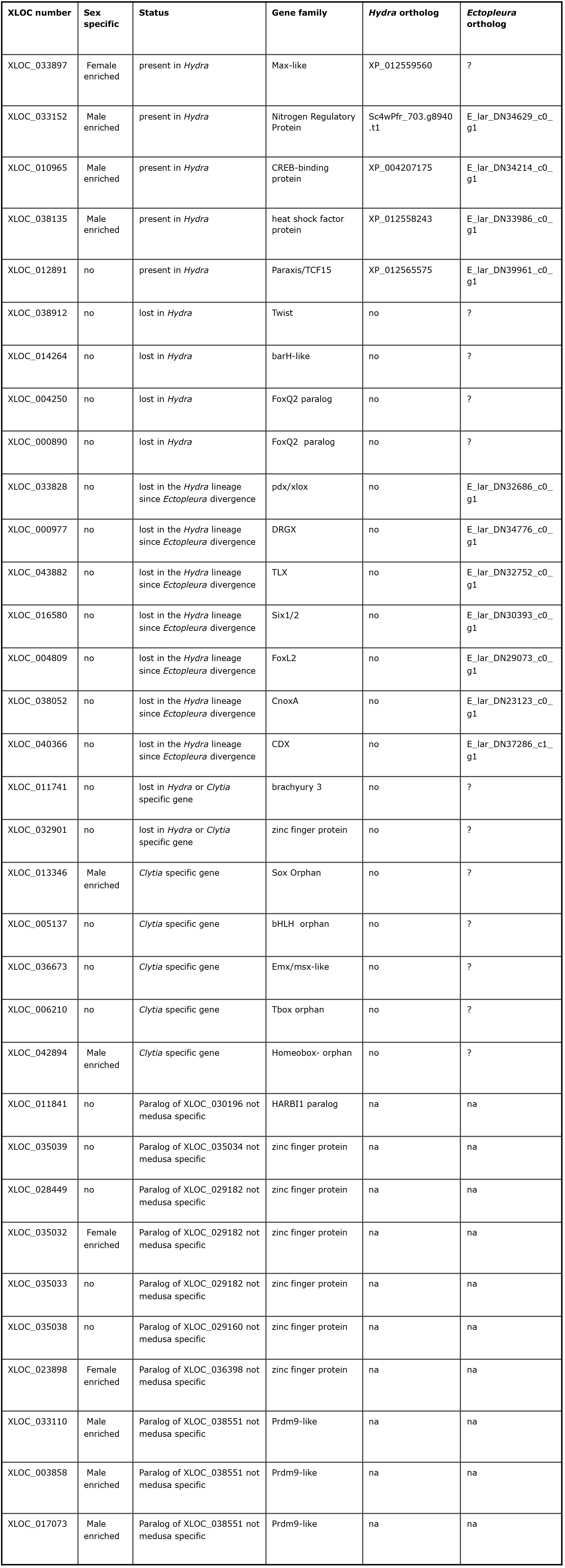
Classification of medusa specific transcription factors. List of *Clytia* medusa specific transcription factor and presence/absence of orthologs in *Hydra* (draft genome) and *Ectopleura* (transcriptome), its closest relative with a medusoid stage. Note that 7 *Clytia* medusa specific TF are absent in *Hydra* but present in *Ectopleura*, suggesting the loss of these TF in the *Hydra* lineage was related to loss of the medusa stage. Ortholog for only one *Clytia* medusa specific TF not sex specific, TCF15, was found in *Hydra*. Na: non-applicable.

**Supplementary Table 5.** Sequencing libraries.

**a) Genomic DNA**
| Library insert size | Technology | Read length | No. of reads |
| --- | --- | --- | --- |
| 180 | Illumina PE | 101 | 688126703 |
| 500 | Illumina PE | 101 | 203497401 |
| 4000 | Illumina Nextra | 101 | 136386432 |
| 8000 | 454 | - | 11802050 |
| 20000 | 454 | - | 9505979 |
| 40000 | 454 | - | 502197 |

**b) RNA preparations for individual libraries**
| Stage | Name | Strain | Nb individuals | Total RNA (ng) | Illumina | cycles |
| --- | --- | --- | --- | --- | --- | --- |
| Early gastrula | EG-1 | Z4B * Z10 | 150 | 1220 | NextSeq | 75 SR |
| Early gastrula | EG-1 | Z4B * Z10 | 150 | 1260 | NextSeq | 75 SR |
| Planula 24hpf | P1-1 | Z4B * Z10 | 150 | 2390 | NextSeq | 75 SR |
| Planula 24hpf | P1-2 | Z4B * Z10 | 150 | 1760 | NextSeq | 75 SR |
| Planula 48hpf | P2-1 | Z4B * Z10 | 150 | 1040 | HiSeq 2500 | 50 SR |
| Planula 48hpf | P2-2 | Z4B * Z10 | 150 | 520 | HiSeq 2500 | 50 SR |
| Planula 72hpf | P3-1 | Z4B * Z10 | 150 | 520 | HiSeq 2500 | 50 SR |
| Planula 72hpf | P3-2 | Z4B * Z10 | 150 | 1105 | HiSeq 2500 | 50 SR |
| Primary polyp | PoPr-1 | Z4B * Z10 | 100 | 494 | HiSeq 2500 | 50 SR |
| Primary polyp | PoPr-2 | Z4B * Z10 | 100 | 450 | HiSeq 2500 | 50 SR |
| Gastrozoid | PH-1 | Z4B | 60 | 224 | HiSeq 2500 | 50 SR |
| Gastrozoid | PH-2 | Z4B | 60 | 936 | HiSeq 2500 | 50 SR |
| Gonozoid | GO-1 | Z4B | 60 | 400 | HiSeq 2500 | 50 SR |
| Gonozoid | GO-2 | Z4B | 60 | 1365 | HiSeq 2500 | 50 SR |
| Stolon | St-1 | Z4B | 60 | 1131 | HiSeq 2500 | 50 SR |
| Stolon | St-2 | Z4B | 60 | 780 | HiSeq 2500 | 50 SR |
| Baby medusa | BMF-R1 | Z4B | 100 | 1523 | HiSeq 2500 | 50 SR |
| Baby medusa | BMF-R2 | Z4B | 100 | 1708 | HiSeq 2500 | 50 SR |
| Mature female medusa | MMF-R1 | Z4B | 1 | 2607 | HiSeq 2500 | 50 SR |
| Mature female medusa | MMF-R2 | Z4B | 1 | 2319 | HiSeq 2500 | 50 SR |
| Mature male medusa | M1 | Z10 | 1 | 1370 | NextSeq | 75 SR |
| Mature male medusa | M2 | Z10 | 1 | 2370 | NextSeq | 75 SR |

**c) Mixed-stage RNA preparation**
| Stage | Strain | individuals | Total RNA (ng) |
| --- | --- | --- | --- |
| Planula P1 | Z4B * Z10 | 150 | 250 |
| Planula P2 | Z4B * Z10 | 150 | 250 |
| Planula P3 | Z4B * Z10 | 150 | 250 |
| Primary Polyp PoPr | Z4B * Z10 | 20 | 250 |
| Gastrozoid/Polyp head PH | Z4B | 60 | 250 |
| Gonozoid GO | Z4B | 60 | 250 |
| Stolon St | Z4B | 60 | 250 |
| Mature Female Medusa MMF | Z4B | 1 | 250 |
| Mature Male Medusa M | Z10 | 1 | 250 |
| **MIX** |  |  | **2250** |

d) Sequences generated for RNA libraries
| Stage |  | Replicate 1 | Replicate 2 |
| --- | --- | --- | --- |
| Early Gastrula | EG | 39331160 | 43845659 |
| One day old Planula | P1 | 38554115 | 42356552 |
| Two day old Planula | P2 | 55900254 | 48155906 |
| Three day old Planula | P3 | 47246549 | 70793282 |
| Stolon | St | 80487282 | 90277892 |
| Primary Polyp | PoPr | 72275162 | 70676083 |
| Polyp Head | PH | 45393098 | 45627654 |
| Gonozoid | GO | 65897403 | 63831768 |
| Baby Medusa Female | BMF | 51835910 | 35779461 |
| Mature Medusa Female | MMF | 44120673 | 49731226 |
| Male Medusa | M | 45793597 | 50284626 |
| Villefranche Mixed | - | 143397155 | - |
| Vienna Medusa | - | 106245738 | - |

**Supplementary Table 6.** Protein datasets used in this study

|  | **download source\*** | **GFF** | **proteins\*\*** |
| --- | --- | --- | --- |
| *Saccharomyces cerevisiae* | www.yeastgenome.org | - | cerpep (13-Jan-2015) |
| *Capsaspora owczazarki* | BROAD origins of multicellularity | - | salpingoeca_rosetta_1_proteins.fasta |
| *Monosiga brevicollis* | BROAD origins of multicellularity | - | monosiga_brevicollis_mx1_1_proteins.fasta |
| *Mnemiopsis leidyi* | https://research.nhgri.nih.gov/mnemiopsis/ | - | ML2.2.aa |
| *Nematostella vectensis* | www.cnidariangenomes.org | nveGenes.vienna130208.nemVec1.gtf | nveGenes.vienna130208.protein.fasta |
| *Amphimedon queenslandica* | NCBI | GCF_000090795.1_v1.0_genomic.gff | GCF_000090795.1_v1.0_protein.faa |
| *Trichoplax adhaerens* | NCBI | GCF_000150275.1_v1.0_genomic.gff | GCF_000150275.1_v1.0_protein.faa |
| *Hydra vulgaris* | NCBI | ref_Hydra_RP_1.0_top_level.gff3 | 102 |
| *Exaiptasia* | NCBI | ref_Aiptasia_genome_1.1_top_level.gff3 | 100 |
| *Orbicella faveolata* | NCBI | ref_ofav_dov_v1_top_level.gff3 | 100 |
| *Acropora digitifera* | NCBI | GCF_000222465.1_Adig_1.1_genomic.gff | GCF_000222465.1_Adig_1.1_protein.faa |
| *Strongylocentrotus purpuratus* | NCBI | ref_Spur_4.2_top_level.gff3 | GCF_000222465.1_Adig_1.1_protein.faa |
| *Saccoglossus kowalevskii* | NCBI | ref_Skow_1.1_top_level.gff3 | 101 |
| *Homo sapiens* | NCBI | ref_GRCh38.p2_top_level.gff3 | 107 |
| *Ciona intestinalis* | NCBI | ref_KH_top_level.gff3 | 102 |
| *Branchiostoma floridae* | NCBI | GCF_000003815.1_Version_2_genomic.gff | GCF_000003815.1_Version_2_protein.faa |
| *Drosophila melanogaster* | NCBI | GCF_000001215.4_Release_6_plus_IS_O1_MT_genomic.gff | GCF_000001215.4_Release_6_plus_IS_O1_MT_protein.faa |
| *Caenorhabditis elegans* | NCBI | GCF_000002985.6_WBcel235_genomic.gff | GCF_000002985.6_WBcel235_protein.faa |
| *Capitella teleta* | NCBI | GCA_000328365.1_Capca1_genomic.gff | GCA_000328365.1_Capca1_protein.faa |
| *Lingula anatina* | NCBI | ref_LinAna1.0_top_level.gff3 | 100 |
| *Crassostrea gigas* | NCBI | ref_oyster_v9_top_level.gff3 | 100 |
| *Aplysia californica* | NCBI | ref_AplCal3.0_top_level.gff3 | 101 |

\* The original Broad institute 'Origins of multicellularity' website is no longer accessible - data downloads are redirected to the NCBI
\*\* For NCBI protein files, an annotation version number is given when the file name is 'protein.faa'

